# Short- and long-term effects of continuous compost amendment on soil microbiome community

**DOI:** 10.1101/2023.02.21.529350

**Authors:** Judith Kraut-Cohen, Avihai Zolti, Nativ Rotbart, Asher Bar-Tal, Yael Laor, Shlomit Medina, Raneen Shawahna, Ibrahim Saadi, Michael Raviv, Stefan J Green, Uri Yermiyahu, Dror Minz

**Author notes:** Equal contribution (Judith Kraut-Cohen, Avihai Zolti). Corresponding author. *Email address* (J. Kraut-Cohen).

## Abstract

Organic amendment, and especially the use of composts, is a well-accepted sustainable agricultural practice. Compost increases soil carbon and microbial biomass, changes enzymatic activity, and enriches soil carbon and nitrogen stocks. However, relatively little is known about the immediate and long-term temporal dynamics of agricultural soil microbial communities following repeated compost applications. Our study was conducted at two field sites: Newe Ya’ar (NY, Mediterranean climate) and Gilat (G, semi-arid climate), both managed organically over 4 years under either conventional fertilization (0, zero compost) or three levels of compost amendment (20, 40 and 60 m^3^/ha or 2, 4, 6 L/m^2^). Microbial community dynamics in the soils was examined by high-and low-time-resolution analyses. Annual community composition in compost-amended soils was significantly affected by compost amendment levels in G (first, second and third years) and in NY (third year). Repeated sampling at high resolution (9–10 times over 1 year) showed that at both sites, compost application initially induced a strong shift in microbial communities, lasting for up to 1 month, followed by a milder response. Compost application significantly elevated alpha diversity at both sites, but differed in the compost–dose correlation effect. We demonstrate higher abundance of taxa putatively involved in organic decomposition and characterized compost-related indicator taxa and a compost-derived core microbiome at both sites. Overall, this study describes temporal changes in the ecology of soil microbiomes in response to compost vs. conventional fertilization.

**Highlights:**

- Dose-dependent changes in soil microbiome structure by manure compost application
- Dynamic short-and long-term changes in soil microbiomes by compost amendment
- Climate, soil properties and management influence compost-amendment effects
- Immediate and temporal cumulative effects of compost on soil α and β diversity

## 1. Introduction

Intensive conventional agricultural practices are largely based on applying mineral fertilizers containing the major elements nitrogen, phosphorus and potassium (NPK) to satisfy crop nutrient requirements and obtain sufficient yields [1]. It is widely understood, however, that the excessive use of chemical fertilizers worldwide has led to extensive environmental pollution [1,2]. A sustainable alternative to mineral fertilization is the use of organic, mainly waste-based fertilizers, such as compost. The use of compost and other organic additives is vital in organic farming and highly recommended to increase soil fertility and health when adopting biodynamic or regenerative practices [3].

Application of organic amendments typically increases the basal soil organic matter content, microbial biomass carbon (C), enzymatic activities and respiration, and results in high N, K, and P availability [3–11]. As organic matter physically stabilizes the soil, its addition also enhances soil structure, mainly reflected in higher aggregate stability, larger aggregate size and higher water retention and lower soil erosion [3,12,13].

The effect and dynamics of long-term organic fertilizer application are more complex. Repeated amendment with compost or manure enriches soil C and N stocks [3], as well as total and available P [2,4,14–17] compared to mineral fertilization [18]. Long-term amendment with compost or manure can either increase or decrease pH, depending on the soil type and its characteristics [3,19]. The slow decomposition of organic fertilizers leads to a prolonged post-application effect on soil biological properties such as microbial biomass C, basal respiration and more [3]. Taken together, these studies support the understanding that application of organic fertilizers over long periods increases soil organic matter and potentially improves soil quality, fertility and structure [2,3].

The soil microbiome is very responsive to environmental cues, which makes it a sensitive marker for evaluating agricultural management [12,20,21]. Several studies have shown that organic amendment increases soil microbial and fungal biomass [2,4,14,22] compared to mineral fertilization. Next-generation sequencing has revealed significant differences between the composition of bacterial and fungal soil communities associated with an organic vs. mineral fertilization regime [2,9,11,19,23]. Twenty years of continuous organic (either compost or manure) vs. conventional farming resulted in significant differences between the respective soil microbial communities [16]. Zhang et al. [19] found a correlation between source soil pH and the effect of fertilization on the soil microbiome. In acidic or near-neutral soils, community composition and Shannon and richness diversity indices were higher with manure vs. mineral fertilization, whereas in alkaline soil, no such effects were observed.

The effect of compost amendment on microbial diversity is complex. According to some studies, bacterial alpha diversity is higher in organically fertilized compared to mineral-fertilized soil [2,9,11,16]. Besides diversity index values, compost also changes the taxonomic composition of microbial communities in agricultural soils. Phylotypes of Alphaproteobacteria, Actinobacteria, Pseudomonas and Bacteroidota were found to increase in denaturing gradient gel electrophoresis profiles under organic fertilization, but only for the short term [24,25], which suggests that at least some of the organic fertilization-related taxonomic changes are transient. These phyla are also highly abundant in composts [26]. In addition, the relative abundances of bacterial taxa known to thrive in nutrient-rich environments, such as Bacillota and Pseudomonadota, were shown to increase in organically fertilized soils [2,16,27]. Conversely, under mineral fertilization, increases in the relative abundance of other bacterial groups, such as Actinomycetota and Chloroflexi, have been documented [2,28].

Most previous studies have focused on one soil type and a single sampling time point after compost amendment, under highly controlled conditions; temporal microbiome changes post-compost have not been examined at high resolution under field conditions. Our study examined short-term (1 week to 2 months) and long-term (annual) temporal microbial dynamics in soils of two field sites: Newe Ya’ar (NY, Mediterranean climate with average winter rainfall of 570 mm per annum), and Gilat (G, semi-arid climate, with average winter rainfall of 238 mm per annum), organically managed over 4 years, using three levels of compost amendment (2, 4 and 6 L/m^2^) vs. conventional fertilization (zero compost). Soils were sampled at low temporal resolution (yearly intervals between 2014 and 2016) and high temporal resolution (weekly to monthly in 2015–2016) post-compost amendment. Our results demonstrate that in semi-arid soil (G site), compost amendment has a long-term dose-dependent effect on microbial community, compared to conventional practice. At the NY site, long-term soil microbiome analysis revealed a cumulative effect of compost amendment, mainly evident in the third year of analysis. Short-term analysis at both sites confirmed that some compost-dependent changes are immediate and visible in the first 4 weeks post-compost amendment: specifically, Shannon diversity was significantly elevated at both sites and the abundance of specific taxonomic groups and operational taxonomic units (OTUs) was altered under compost vs. conventional fertilization.

## 2. Materials and methods

### 2.1. Site description and sampling

The experiments were conducted at two sites: Newe Ya’ar Research Center (NY) in northern Israel (32°70′N, 35°18′E), characterized by a Mediterranean climate where the annual winter rainfall is 570 mm, and soil type is a clay soil (64% clay,18% silt, 18% sand), Vertisol-type, classified as Chromic Haploxerert; and Gilat Research Center (G) located in southern Israel (31°20’N, 34°41’E), and characterized by a semi-arid climate, with annual winter rainfall of 238 mm. The soil type at G is sandy loam (25% clay, 35% silt, 40% sand), Loess, classified as Calcic Xerosol. Soil parameters for both sites are described in Rotbart et al. [29].

Both G and NY sites have been organically managed since 2011, using three levels of compost amendment (20, 40 and 60 m^3^/ha, treatments of 2, 4, 6 L/m^2^, respectively; referred to herein as treatments 2, 4, 6, respectively) or conventional fertilization (zero compost; treatment 0). Aside from fertilization, all treatments, including the conventional fertilization, were managed under organic practices with respect to plant protection and in NY, green manure was incorporated as well. Soils were sampled as described in section 2.2. Briefly, DNA was extracted and 16S rRNA gene amplicons were sequenced for microbiome analysis. The experimental timeline is shown in **Fig. 1**.

**Fig. 1.**
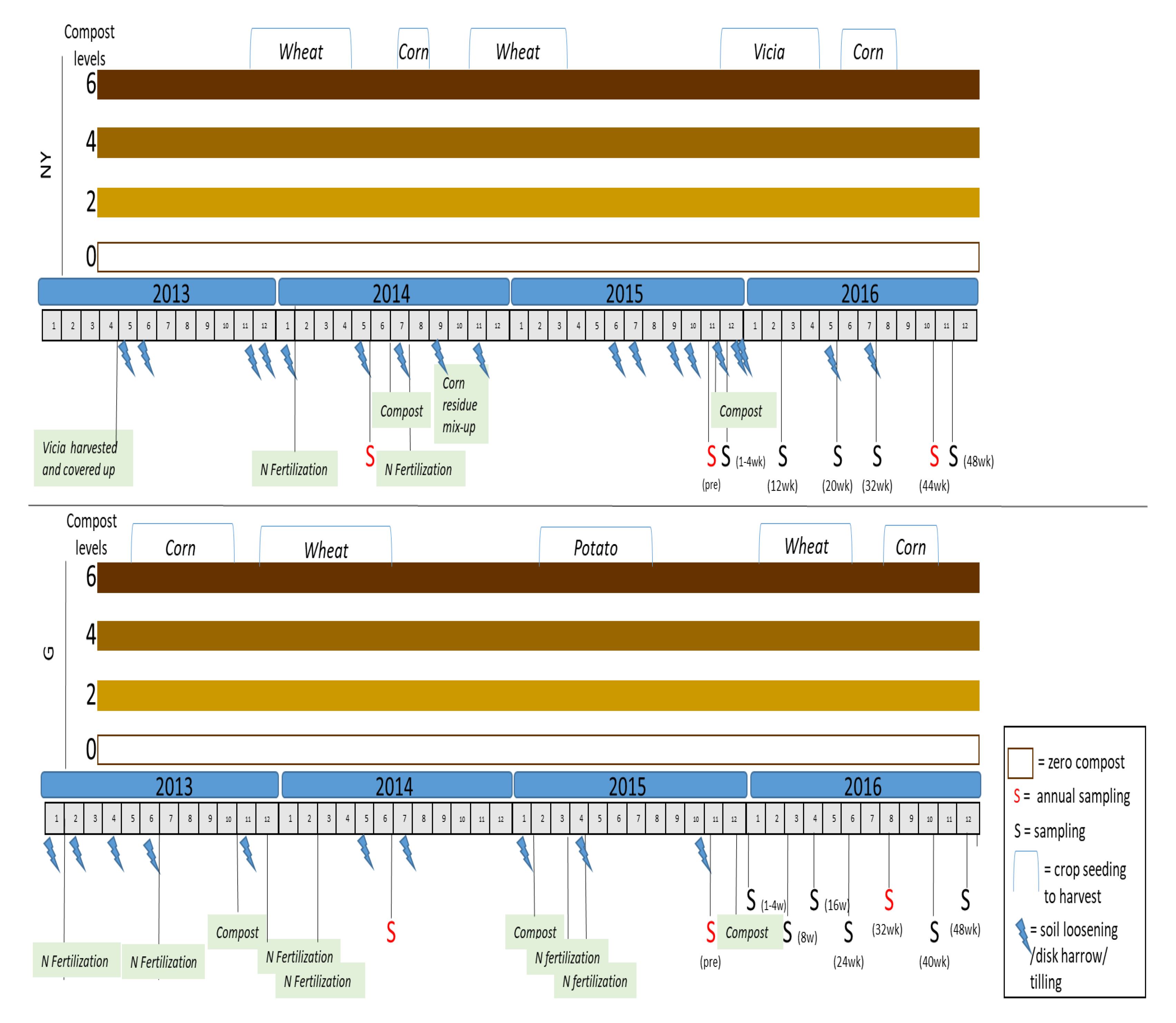
Experimental timeline in Gilat (G) and Newe Ya’ar (NY) sites. The experiment was conducted between 2013 and 2016. Each field site (G, NY) had four treatments: 0 (conventional fertilization) and 2, 4, 6 (20, 40 and 60 m^3^/ha of compost amendment, respectively). The crops grown at each site are indicated at the top. All sampling events are marked with "S": a red "S" indicates that the sampling event was part of the annual analysis from 2014 to 2016. Lightening symbols represent soil cultivation (tillage/loosening).

Section 2.3 and **Table S1** summarize the main properties of the composts applied during the experimental period. The compost was organic cattle manure-based and applied at levels of 2, 4 and 6 L/m^2^, equivalent to 20, 40 and 60 m^3^/ha, respectively (resulting in 10.9, 21.9 and 32.8 t/ha in G and 8.7, 17.4 and 26.1 t/ha in NY, respectively). The resulting applied quantities of organic C were 2.1, 4.1 and 6.2 t/ha in G and 2.4, 4.8 and 7.3 t/ha in NY, respectively. In organic farming, 20 to 40 m^3^/ha are recommended annual treatment doses, and 60 m^3^/ha per year was selected as two times the median value between 20 and 40. The conventional plots of "no compost" (0 treatment) were fertilized with urea or ammonium sulfate. In both organic and conventional treatments at NY, all plots were amended with harvested vetch (*Vicia sativa*), a commonly used green manure in agriculture. Each of the sites (G, NY) included 3–5 replicates of the four treatments (0, 2, 4, 6). The treatment plots were arranged in 3–5 blocks, each block containing the four treatment plots (0, 2, 4, 6) in random order. All plots have had field crops in rotation since 2009. All plots, including the conventional-practice plots, were managed without pesticide use. The experiment in the current settings was initiated in 2010 and sampling for the microbiome measurements was begun in 2014 (see section 2.2) [29,30]. Rainfall records (in mm) during the experimental period are presented in **Fig. S1**.

#### 2.1.1. Fertilization in G

Compost-treated plots in G were amended with compost in October 2013, February 2015 and December 2015. Urea fertilizer was applied on the zero compost plot in June 2013, December 2013 and February 2014, and ammonium sulfate (21% N) fertilization was applied in March 2015, April 2015 and May 2015.

Shallow soil cultivation was applied in all plots (including the conventional-practice treatment) using a disc harrow. All field treatments, crop rotations and sampling points are illustrated along the experimental timeline in **Fig. 1**. Compost chemical properties are detailed in **Table S1**.

#### 2.1.2. Fertilization in NY

In April 2013, all NY plots were amended with harvested vetch (*Vicia sativa*) as green manure. Chemical fertilization on the zero compost plot was applied in January 2014 (potassium chloride (KCl) and 11% P) and with urea (46% N) in July 2014. Because the legume vetch was cultured from November 2015 to April 2016, no additional N fertilization was added in that growing season. Compost-treatment plots were amended with compost in June 2014 and November 2015, according to the specified compost levels (**Fig. 1**).

### 2.2. Sampling

The overall area of each field site was 1 ha, which was subdivided into 20 plots, and the dimension of each plot was 18 m x 30 m = 540 m^2^. Soil samples were collected at high frequency (weekly to monthly) for 1 year and at low frequency (once annually) for 3 years. Annual sampling was conducted yearly between 2014 and 2016 in June 2014, November 2015 and August 2016 (G), and in May 2014, November 2015 and October 2016 (NY). High-resolution sampling was conducted in 2015 and 2016 on the following dates: G – pre-compost (November 2015), and 1, 2, 3, 4, 8, 16, 24, 32, 40 and 48 weeks post-compost application; NY – pre-compost (November 2015) and 1, 2, 3, 4, 12, 20, 32, 44 and 48 weeks post-compost application. At each sampling, a soil composite sample was taken from 0–15 cm depth. Soil samples were cleaned of stones and plant material, and kept at −80 °C for DNA extraction. In contrast to laboratory experiments in a controlled setting, field soils are highly heterogeneous. Therefore, care was taken to ensure that the sampling would provide an as accurate as possible representation of the site soils. Soil was sampled from five blocks, composed of randomized plots, with each sample composed of three separate soil samples that were pooled before the final DNA extraction.

### 2.3. Analysis of compost and soil chemical properties

Commercial compost (NY – Compost Sade Eliyahu–Agricultural Cooperative Society Ltd., G – Machluf, Northern Negev manufacturers Hayun Ecology Compost Industry–Cooperative Society Ltd.) consisting of cattle manure was applied to each of the organic plots. Organic matter (OM) contents of the compost varied between 24.0 and 53.9%, organic C between 14.1 and 31.7%, total N (TN) between 1.03 and 1.9% and C:N ratio between 12.6 and 16.8. Chemical and physical characteristics of the applied compost during the experiment are described in **Table S1**. Soil for the analysis of chemical properties (presented in **Table S2)** was sampled from layer 0–15 cm, at the following time points: May 2014, May 2015, and November 2016 in NY and June 2014, June 2015 and November 2016 in G. The soil samples were brought to the laboratory, dried in an oven at 40 ℃, ground, and sieved through 2 mm sieves. Total organic carbon (TOC) and TN were determined using a C/N analyzer (Flash EA 1112, Thermo Fisher Scientific). Soil texture was determined using the hydrometer method based on Stokes’ law (analyzed by Dr. G.J. Levy; partly presented in Sharma et al. [30]). Water extracts at a 1:5 soil:water ratio (w/v) were obtained, filtered through a slow filter paper (#293; 1–2 µm Sartorius, Thermo Fisher Scientific), and then used to determine electrical conductivity (EC), pH and water-soluble ions. The EC and pH were determined with respective meters. Water-soluble sodium ions (Na^+^) and K^+^ were determined in the extract with a flame photometer (Sherwood Scientific Ltd., Cambridge, UK), and the concentration of Cl^-^ was determined using a Discrete Auto Analyzer (Gallery plus, Thermo Fisher Scientific). Available nitrate-N (NO_3_-N) and ammonium-N (NH_4_-N) concentrations were determined by soil extraction with 1 mol/L KCl, and available P by soil extraction with 0.5 mol/L sodium bicarbonate (NaHCO_3_). The extracted NO_3_- N, NH_4_-N and P were measured calorimetrically using the Discrete Auto Analyzer. In NY, TOC and TN were measured only in 2014–2015. Na in the soil solution was determined only in G samples.

The main effects of compost level, blocks, sampling events and their interactions on measured variables were determined statistically using the three-way ANOVA procedure of JMP 16. The significance of comparisons among treatments was tested by Tukey–Kramer honest significant difference (HSD) test at *P* < 0.05.

### 2.4. DNA extraction and amplicon sequencing

For analysis of community composition at each site, five replicate samples from each treatment were analyzed for the years 2015 and 2016, and three replicates for 2014. In total, 232 (G) and 212 (NY) soil samples were collected over a 3-year period. Total genomic DNA was extracted from 0.1 g of soil using the Exgene soil DNA isolation kit (GeneAll Biotechnology Co. Ltd., Republic of South Korea) as previously described [21]. DNA was PCR-amplified according to Earth Microbiome Project recommendations, using primer pair 515F and 806R targeting the variable V4 region of microbial small subunit rRNA genes [31] using a two-stage protocol, as described previously [32]. Amplicons from the first-stage PCR were used as templates for the second-stage PCR employing Access Array Barcode Library for Illumina Sequencers primers (Fluidigm, San Francisco, CA). Libraries were pooled and sequenced, with a 20% phiX spike-in, on an Illumina MiSeq employing V3 chemistry. Fluidigm sequencing primers targeting the CS1 and CS2 linker regions were used to initiate sequencing. De-multiplexing of reads was performed on-instrument. Library preparation and pooling were performed at the University of Illinois at Chicago Genomics Research Core. MiSeq sequencing was performed at the W.M. Keck Center for Comparative and Functional Genomics at the University of Illinois at Urbana-Champaign. Sequences were uploaded to NCBI under Bioproject PRJNA688413.

### 2.5. Sequence analysis

Samples were sequenced in two batches: 2014 samples, and then all 2015–2016 samples. Sequences were analyzed at yearly resolution as well as at high resolution between 2015 and 2016, as described in section 2.2. In the 2014–2016 annual analysis, we analyzed 897,128 and 886,079 high-quality sequences originating from 50 (NY) or 51 (G) samples, which clustered to 4128 OTUs. The minimum number of sequences per sample was 11,077 (G) and 11,239 (NY), and the maximum number of sequences per sample was 19,497 (G) and 20,178 (NY), with a mean of 18,990 (G) and 17,942 (NY) sequences per sample.

In the high-resolution analysis, 17,605,266 sequences originating from 410 samples were generated and analyzed. These sequences were clustered into 9373 (NY) and 5924 (G) OTUs. The minimum number of sequences per sample was 13,037, the maximum was 14,525 and the median number of sequences per sample was 13,714. The bioinformatics pipeline was as described previously [33]. Briefly, forward and reverse paired-end reads were joined, filtered by quality and trimmed using the MOTHUR software package [34]. Clustering sequences into *de novo* OTUs by 97% similarity, and classification and depiction of differentially abundant taxa were performed with QIIME version 1.9 [35], using Silva 123 as the reference database [36]. Chimeric sequences were removed using UCHIME [37]. Chloroplast and mitochondrial sequences were also removed. All generated OTU tables were rarified to the fewest number of sequences per sample before alpha diversity was calculated. All OTUs with fewer than 50 sequences per OTU in all datasets were removed from the dataset. Three samples (two from NY, November 2015, 40 m^3^/ha compost treatment and one from G, August 2016, 20 m^3^/ha compost treatment) failed to generate sufficient reads and were removed from the dataset. Three to five biological replicates per treatment were analyzed on each date. The high-resolution OTU BIOM and taxonomy files were exported into QIIME 2 [38] and analyzed using the bioinformatics tools described in sections 2.6–2.9.

### 2.6. Diversity analysis

The compositional dissimilarity between samples (beta diversity) throughout 2016 was measured using the Bray–Curtis distance matrix and visualized using non-metric multidimensional scaling (nMDS) plots [39] (annual resolution) or principal coordinate analysis (PCoA) (high temporal resolution).

PCoA for the soil treatments (0, 2, 4, 6) was created based on QIIME 2 Bray– Curtis dissimilarity index data, and plotted by ggplot2 package in R [38] with centroids representing 95% normal data confidence ellipses for each group.

Pairwise comparisons between average Bray–Curtis distance matrices were calculated for compost treatments compared to conventional practice at each time point, and plotted using the ggplot2 package in R. Significance of dissimilarity was estimated by Kruskal–Wallis non-parametric test (significance threshold of *P* < 0.05); pairwise groups within each category (year, treatment) were compared using analysis of similarities (ANOSIM) test and treatment pairs in each year were compared using permutational multivariate ANOVA (PERMANOVA) test.

Feature tables were rarified to the minimal feature number per sample (in G 13,000 and in NY 8500 for 2016, and 11,000 for both NY and G in the 3-year analysis) before Shannon diversity indices were calculated using QIIME 2. Taxonomic differences were calculated by Mann–Whitney test (high-resolution analysis, all time points) or Wilcoxon test (yearly analysis) and considered significant at *P* < 0.05.

### 2.7. Heat map

For the high-resolution analysis in 2016, all significantly (Kruskal–Wallis test, FDR-corrected *P* < 0.05) enriched or depleted OTUs from each of the compost treatments as compared to the conventional-practice treatment were plotted. Significantly different OTUs between treatments 0 and 6 were selected and plotted for all treatments (0, 2, 4, 6) on the heat map, along the full experimental timeline (either yearly or weekly–monthly resolution), as described in section 2.2. The differentially abundant OTUs were sorted taxonomically at the phylum level, as well as by abundance patterns in each treatment. The heat maps were created using the ‘pheatmap’ package in R [40].

### 2.8. Core microbiome analysis

The core microbiome of all compost treatments together (all compost samples were combined) and the core microbiome (set at 100% prevalence) of the conventional treatment (0) were computed individually for G and NY using the core features plugin in QIIME 2 with at least 100% prevalence. Venn diagrams were created at the OTU level to compare different core microbiomes using Venny [41].

### 2.9. ANCOM and MaAsLin2 analysis

Identification of significantly differentially abundant OTUs between treatments (0, 2, 4, 6) was determined by analysis of composition of microbes (ANCOM) among all treatments using the ANCOM plug-in in QIIME 2 [42], with FDR-corrected *P*-values already embedded in the ANCOM test. In addition, centered log ratio-transformed data were used to identify differentially abundant features between the compost and conventional treatments with the Microbiome Multivariable Associations with Linear Models (MaAsLin2) package in R [43]. Significant OTUs derived from both methods were compared by Venny [41], and OTUs that were found significant in both were further plotted using the ‘pheatmap’ package in R [40].

## 3. Results

The short-and long-term effects of compost vs. conventional fertilization on microbial community composition were examined over time, at sites G and NY that differ in their soil type [29], climate and crop rotation (detailed in section 2.1).

### 3.1. Effect of yearly compost amendment vs. conventional practice on soil chemistry and community composition

In the semi-arid soil (G), all measured chemical parameters except NH_4_, Cl and pH were significantly elevated in a compost dose-dependent manner (**Table S2**). In the Mediterranean soil (NY), TN, P and K were significantly elevated in compost treatments (in a dose-dependent manner for TN and P) while NO_3_ exhibited a similar trend. The effect of compost amendment (compost vs. conventional practice) and dose of the amended compost (treatments 2, 4, 6) on microbial community composition was examined yearly. Soils were sampled at site G in June 2014 (Y1), November 2015 (Y2) and August 2016 (Y3) and at site NY in May 2014 (Y1), November 2015 (Y2) and October 2016 (Y3).

The most dominant phyla in the full dataset (all annual sampling times and treatments) were Pseudomonadota (27% in NY, 28% in G), Actinomycetota (22% in NY, 21% in G), Bacillota (15% in NY, 13% in G), Thaumarchaeota (12.8% in NY, 8.1% in G) and Bacteroidota (4% in NY, 8.8% in G).

The effect of compost amendment at annual resolution was initially examined by pairwise comparison based on the Bray–Curtis similarity indices of the three compost levels vs. the conventional treatment. A significant difference for the "year" (*P* < 0.0001 in G and NY) and "compost" (*P* =0.007, in G) parameters was found by ANOSIM for all 3 years (2014–2016). However, the "compost" parameter was not significant in NY samples.

At both sites, all tested years (Y1, Y2, Y3) varied significantly from each other (**Table S3 A** and **B**). These differences represent dynamics that may reflect the variation between years in soil management, crop, weather and sequencing batch.

Significant shifts in G soil microbial community structure were observed at the higher rates of compost amendment (4, 6) relative to conventionally treated soil (0) across all 3 years of the study. Conversely, at the NY site, no significant changes were found between treatments pairs when comparing all 3 years together (**Table S3 C** and **D**).

The difference in microbial community structure of soils receiving compost and those receiving conventional treatment was visualized using box plots based on dissimilarity comparison of Bray–Curtis distances for each year (**Fig. 2 A and B**). Comparison of compost treatments revealed significant changes. In G, comparison of compost treatments within each year revealed significant and cumulative changes between treatments: treatments 2 and 6 in Y1, between 2 and 6, and 4 and 6 in Y2 and between 2 and 6, 4 and 6, and 2 and 4 in Y3 (**Fig. 2 A, Table S3 E**). In NY, significant differences were found between treatments 2 vs. 6, and 4 vs. 6 in Y3 only (**Fig. 2 B, Table S3 F**).

**Fig. 2.**
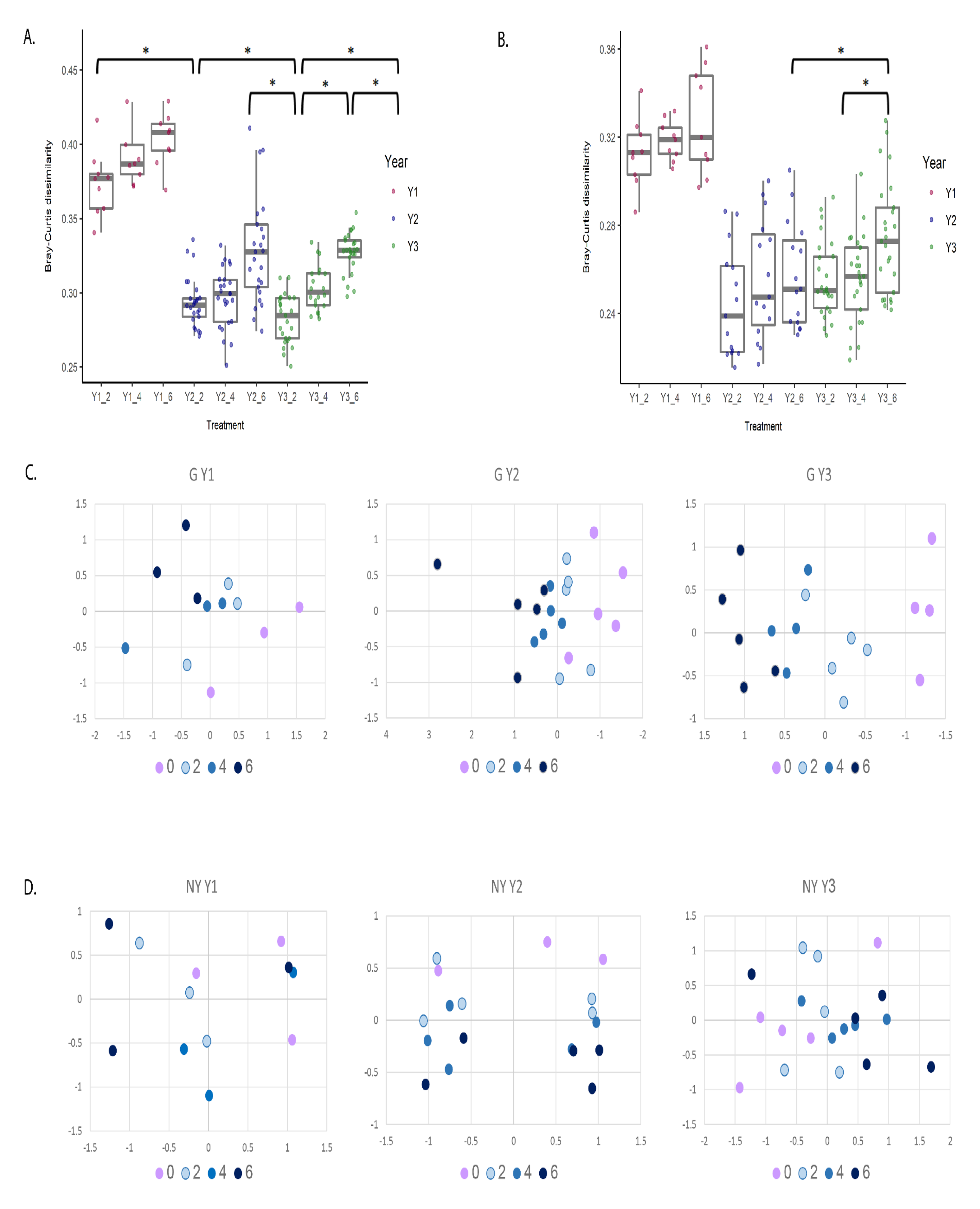
Compost amendment alters soil community composition. Bray–Curtis dissimilarities between microbiomes of compost-amended soils (2, 4, 6) and conventionally fertilized soil (0) for the period 2014–2016 at yearly resolution. Pairwise comparisons are indicated for soils from G (A) and NY (B). Asterisks represent significant differences between groups (*P* ≤ 0.05, Wilcoxon test). The 2014 (Y1) samples are marked in red, 2015 (Y2) in blue and 2016 (Y3) in green. nMDS ordination was plotted for the Bray–Curtis dissimilarities in soil microbial community composition (OTU level) at G (C) and NY (D) sites between 2014 and 2016. Conventionally fertilized (0) samples are marked in violet, and compost-amendment treatments are marked in light blue (2), blue (4), and dark blue (6). Stress values were <0.1 for all analyses.

Samples were visualized using nMDS (**Fig. 2 C** and **D**), and statistical analyses (PERMANOVA) were conducted to determine whether differences in microbial community structure between treatments was significant (**Table S4**). A cumulative microbiome response to compost levels in G soils was visualized using nMDS (**Fig. 2 C**). In Y1 at the G site, the difference between the different fertilization treatment groups was not significant (**Fig. 2 C**, **Table S4**). In Y2 and Y3, bacterial communities in soil amended with the lowest compost level (treatment 2) differed significantly from that with the highest compost level (treatment 6) (**Table S4**). Conventionally fertilized soil (treatment 0) differed significantly from the compost treatments (4 and 6 in Y2 and 2, 4 and 6 in Y3) (**Fig. 2 C**, **Table S4**).

At NY, nMDS analysis revealed a fertilization and dose-dependent trend that was not significant due to the high variation between replicates in microbial profiles (**Fig. 2 D**, **Table S4**). No significant differences were found in the alpha diversity indices between the treatments at either site (not shown).

### 3.2. Effect of annual compost amendment vs. conventional practice on microbial community structure

Several differentially abundant taxonomic groups were characterized in G soils during the 3 years of observation. In Y1 no differences were found. In Y2, the relative abundance of OTUs from bacterial orders Gemmatimonadales and Solirubrobacterales was lower in compost-amended soils compared to conventional fertilization (*P* = 0.0217 and 0.034, respectively, Wilcoxon test). However, none of the 17 tested orders were significant after false-discovery rate correction (i.e., FDR q-value) (**Table S5**). In Y3, the relative abundance of Rhizobiales was significantly lower in the compost vs. conventional-practice soils. In addition, compost level-dependent patterns were found in the relative abundance of the taxa Cytophagales and Myxococcales, usually associated with high organic matter consumption, using Wilcoxon test to compare different doses of compost treatment vs. conventional practice in Y3 (**Table S5**). In G soils, the relative abundance of 134 OTUs differed significantly between conventional fertilization and the high-compost treatment (0 vs. 6), whereas the relative abundance of only 4 OTUs was significantly different between 0 and 4 soils. No significant difference was found between the other treatments. For the most part, the relative abundance of OTUs in the phyla Bacillota and Bacteroidota was greater in compost-amended soils, whereas OTUs from the Euryarchaeota showed higher relative abundance in the conventional-practice soil (**Fig. S2**). We did not detect any OTUs that differed significantly between treatments at the NY site in the annual analysis.

Overall, unlike in NY, compost amendment affected microbial community composition in the semi-arid G soils. After 3 years of analysis of an ongoing experiment, we could detect a clear and cumulative influence of compost amendment, suggesting a long-lasting effect.

### 3.3. High-resolution microbial community temporal dynamics throughout 2016

To further understand the temporal dynamics of bacterial communities in response to compost amendment, we sampled both G and NY soils throughout 2016 at high temporal resolution.

During this high-resolution sampling year, the dynamics of compost amendment were examined at both sites. The dominant phyla in the high-resolution analysis were Pseudomonadota (28% in G and 27% in NY) and Actinomycetota (20% in G and 26% in NY). In G, there was a clear gradient between treatments across all sampling time points, and PERMANOVA demonstrated significant differences between all pairwise treatment comparisons (0, 2, 4, 6) (**Fig. 3 A**, **Table S6**). At the NY site, treatment groups were less strongly clustered, but a similar trend was observed, with significant differences between compost level treatments (2, 4, 6) and the conventional treatment (0), and between low (2) and high (6) compost treatments (**Fig. 3 B, Table S6**).

**Fig. 3.**
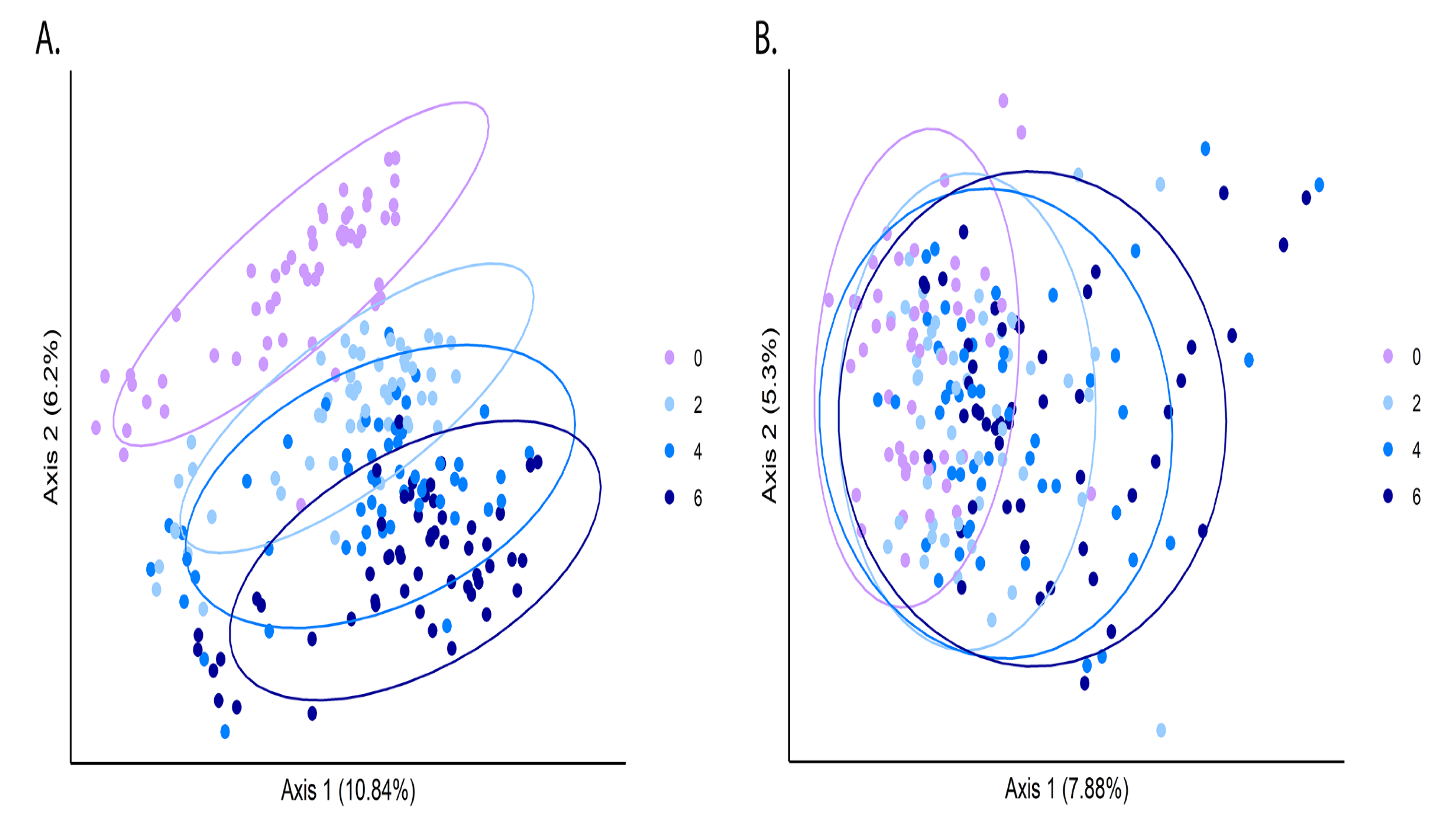
High-resolution analysis of compost-mediated changes in soil microbiomes. Ordination of soil microbiomes was conducted using PCoA based on a Bray–Curtis dissimilarity matrixof all high-resolution-sampled time points in 2016, in G (A) and NY (B) soils. Conventionally fertilized (0) samples are marked in violet, and compost-amendment treatments are marked in light blue (2), blue (4), and dark blue (6).

At both sites, agricultural management and environmental conditions caused dynamic temporal changes in community structure. To focus on the effects of compost amendment for each sample and time point, we plotted Bray–Curtis differences compared to the site’s conventional-practice control (0) at the same time point (**Fig. 4 A** and **B**). Compost, at any level of amendment, induced an immediate and sharp shift in microbial community structure, followed by a dynamic change in the community (**Fig. 4 C** and **D**). Starting at 12 weeks post-compost amendment, the community dynamics stabilized. Nevertheless, some residual effects of compost amendment on soil microbial communities lasted 32 or 44 weeks (Y3 in **Fig. 2 A** and **B**) and 1 year (**Fig. 4 A** and **B**), demonstrating that compost significantly impacted the structure of soil microbial communities over the long term. In G, comparing the distance in community composition at each sampling time to the equivalent soil community in the conventional practice, a significant dose-dependent response to the compost was observed immediately after compost application. This effect continued throughout the high-resolution sampling period and was still detected 1 year after compost amendment (**Fig. 4 A**; Mann–Whitney pair test *P*-values: 0 vs. 2, *P* = 0.00008; 0 vs. 4, *P* = 0.035; 0 vs. 6, *P* = 0.00018). In NY, moderate differences between treatments were observed, and a significant effect was found only between conventional practice and the high compost treatment in the high-resolution sampling set (Mann–Whitney 0 vs. 6, *P* < 0.005) (**Fig. 2 D**, **Table S4** and **Fig. 4 B**). At the OTU level, a total of 1290 (G) and 540 (NY) OTUs differed significantly (Kruskal–Wallis test, *P* < 0.05) in abundance between the compost treatment and conventional fertilization samples (**Fig. 4C** and **D**, **Fig. S3**). At both sites, an immediate response to compost amendment was observed at all compost levels. In addition to this immediate shift in abundance, we also noted a dynamic set of changes in communities during the weeks after compost was applied (**Fig. 4 C** and **D**, **Fig. S3**). At both G and NY sites, OTUs assigned to the phyla Bacteroidota, Bacillota and Pseudomonadota were tightly and positively associated with compost amendment. Gemmatimonadetes, and Verrucomicrobia showed a mixed response at both sites, with some OTUs increasing and others decreasing in relative abundance in response to compost amendment (**Fig. 4 C** and **D**).

**Fig. 4.**
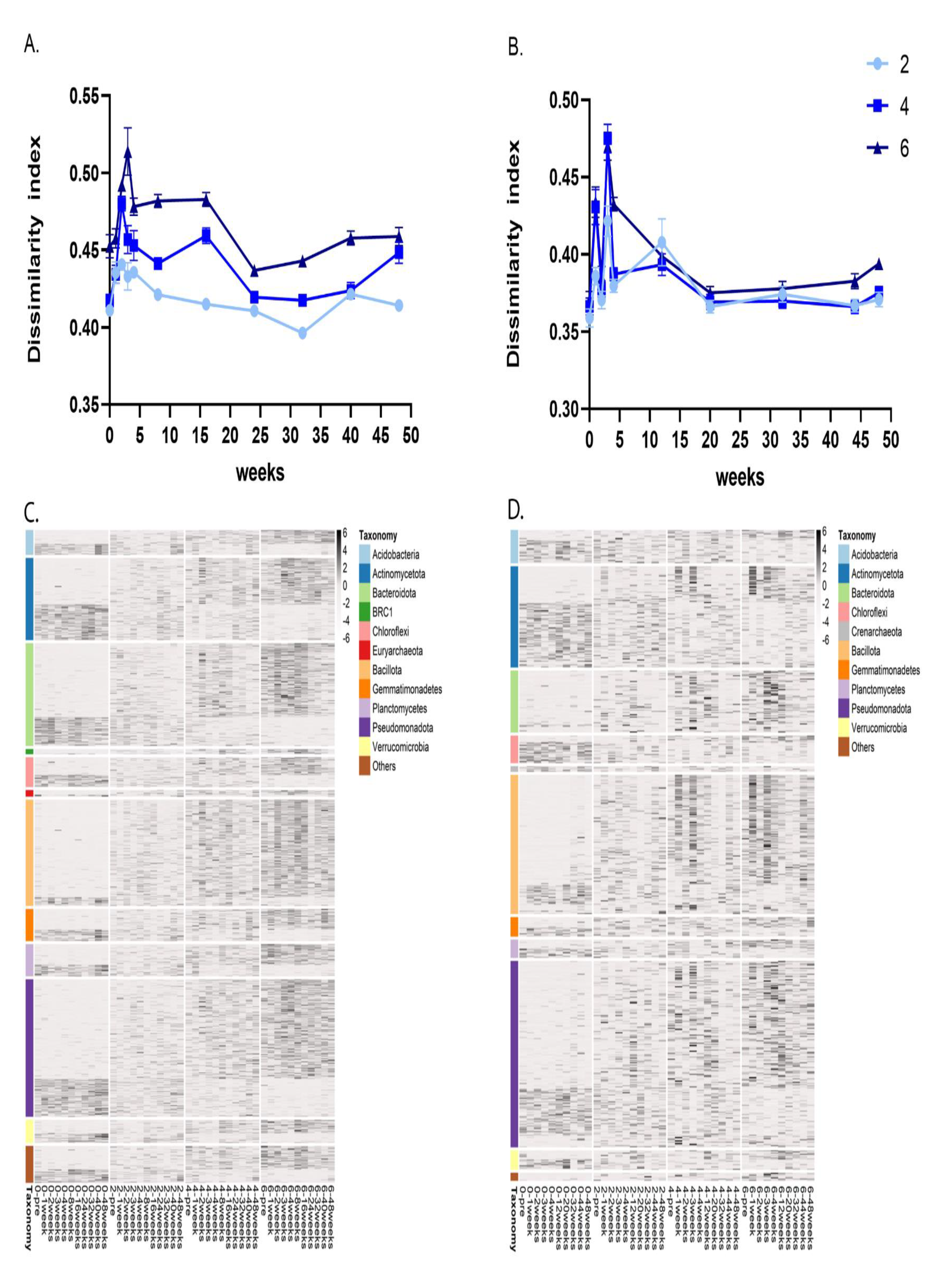
High-resolution temporal dynamics of compost-mediated shifts in microbial community structure during 2016. Bray–Curtis dissimilarities between microbiomes of the compost-amended soils—treatments 2, 4, 6 (marked with light blue, blue and dark blue, respectively)—and that of conventionally fertilized samples (treatment 0) at high temporal resolution over 2016 were visualized for G (A) and NY (B) soils. Heat maps of treatment-enriched OTUs at multiple sampling points in 2016 are shown for G (C) and NY (D) sites. All visualized OTUs were significantly (Kruskal–Wallis test, FDR-corrected *P* < 0.05) enriched or depleted in treatment 0 compared to any of the compost treatments (2, 4 or 6). The heat map presents average OTU abundance (for all corresponding replicated treatments) at each sampling point, normalized per row (Z-score). The colored annotation bars represent the classification of each OTU at the taxonomic level of phylum.

When the mean average of the relative abundance of each phylum in this analysis was plotted over time, we observed an increase in the relative abundance of some phyla in response to compost amendment: Bacillota, Pseudomonadota and Bacteroidota at both sites (**Fig. S3 A** and **B**), and *BRC1*, Planctomyces and Acidobacteria in G **(Fig. S3 A)**. The relative abundance of other phyla increased over time in the conventional-practice (0 compost) samples, especially in NY. These phyla included Actinomycetota and Chloroflexi, and for early time points (up to week 20), also Acidobacteria and Crenarchaeota **(Fig. S3 A)**. These observations were likely due to the dynamics of organic matter degradation and composition with time post-amendment. Thus, the relative abundance of many microbial taxa was altered in a dose-dependent fashion in response to compost amendment. A clear gradient between abundance patterns for levels 2, 4 and 6 of compost treatment was observed (**Fig. 4 C** and **D**, **Fig. S2**).

### 3.4. Compost amendment increases soil community diversity

Next, we compared microbial community diversity by comparing Shannon diversity indices for all treatments in 2016 (high temporal resolution) after rarefaction. The initial diversity in G soil was higher than that of NY (Shannon index average was 10.86 in G, 10.22 in NY, for all treatments at a rarefied sequence depth of 13,000 in G and 8500 in NY sequences/sample). Compost amendment increased alpha diversity relative to conventional practice at both sites (**Fig. 5**). At NY, higher compost levels resulted mostly in higher diversity, and all compost treatments were significantly different from the conventional practice across the dataset (Kruskal–Wallis pair test; treatments 2, 4, 6, *P* < 0.0001). Alpha diversity levels were also significantly different between the highest and lowest compost treatments (2 vs. 6; *P* < 0.08) (**Fig. 5 B**). At G, alpha diversity (Shannon index) was significantly higher, and similar to one another, in all compost treatment samples relative to conventional-practice control samples (Kruskal–Wallis pair test; treatments 2, 4, 6, *P* < 0.001; **Fig. 5 A**).

**Fig. 5.**
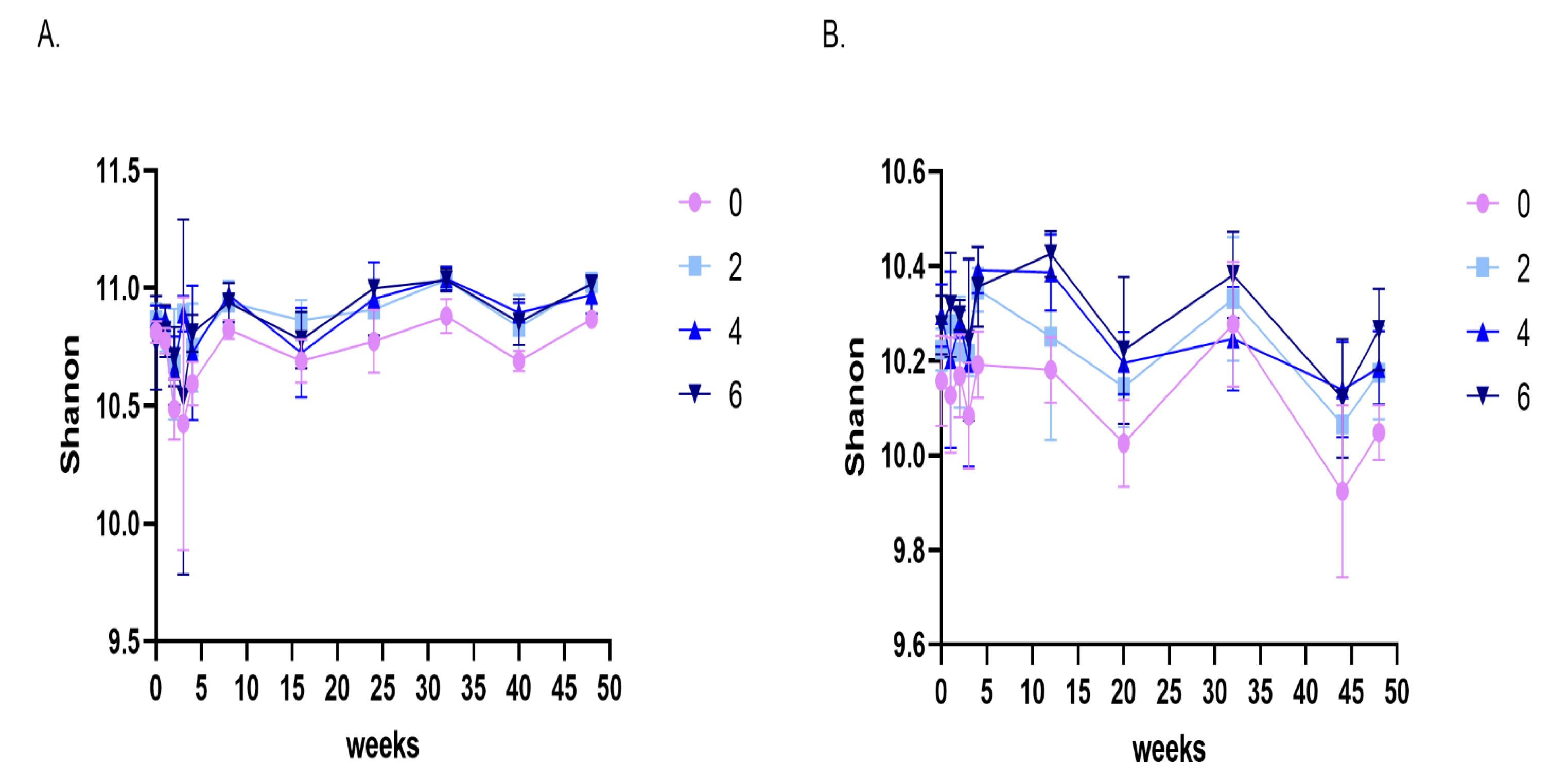
Temporal dynamics of alpha diversity in the microbiomes of compost-amended and conventionally fertilized soils during 2016. Alpha diversity, as measured by Shannon diversity index, in conventional-practice (0) samples is marked in violet and compost-amendment treatments in light blue (2), blue (4), and dark blue (6), from the high-resolution 2016 sampling in G (A), and NY (B).

### 3.5. Compost-amended soils have unique microbial populations

We identified 332 core OTUs in conventionally fertilized soils vs. 205 OTUs in compost-amended soils at the G site. Similarly, we identified 268 core OTUs in conventionally fertilized soils vs. 174 OTUs in compost-amended soils at the NY site. At each site, the core microbiome was determined based on OTUs represented in all compost-amended soil samples from all sampling time points (treatments 2, 4, 6). This site-specific core microbiome was compared to that of the conventional treatment. In G, 27 core OTUs were unique to the compost-amended soils, and 121 were unique to the conventionally fertilized soils (**Fig. S4 A**). In NY, 7 OTUs were unique to compost-amended soils and 80 OTUs were unique to the conventional treatment (**Fig. S4 B**). In G, most compost core OTUs were annotated to Bacillota (9) and Pseudomonadota (7), while in NY, most compost core OTUs were annotated to Actinomycetota (5) and Pseudomonadota (4). Most conventional-treatment core OTUs at both sites were annotated as Pseudomonadota (37 in G and 31 in NY) and Actinomycetota (32 in G and 28 in NY). Alphaproteobacteria (26 in G and 20 in NY), Actinobacteria (16 in G) and Thermoleophilia (13 in NY) were the most abundant class-level annotations in conventional-treatment core OTUs. Bacilli (9) in G, and Gammaproteobacteria at both sites (4 in G, 3 in NY) were the most abundant class-level annotations in the compost core OTUs. When the core communities from both experimental sites were compared, we found 2 OTUs that were shared by the compost-amended soils of G and NY; 12 OTUs were shared by the core communities of the conventionally fertilized soils at both sites (**Fig. 6 A, Table S7**).

**Fig. 6.**
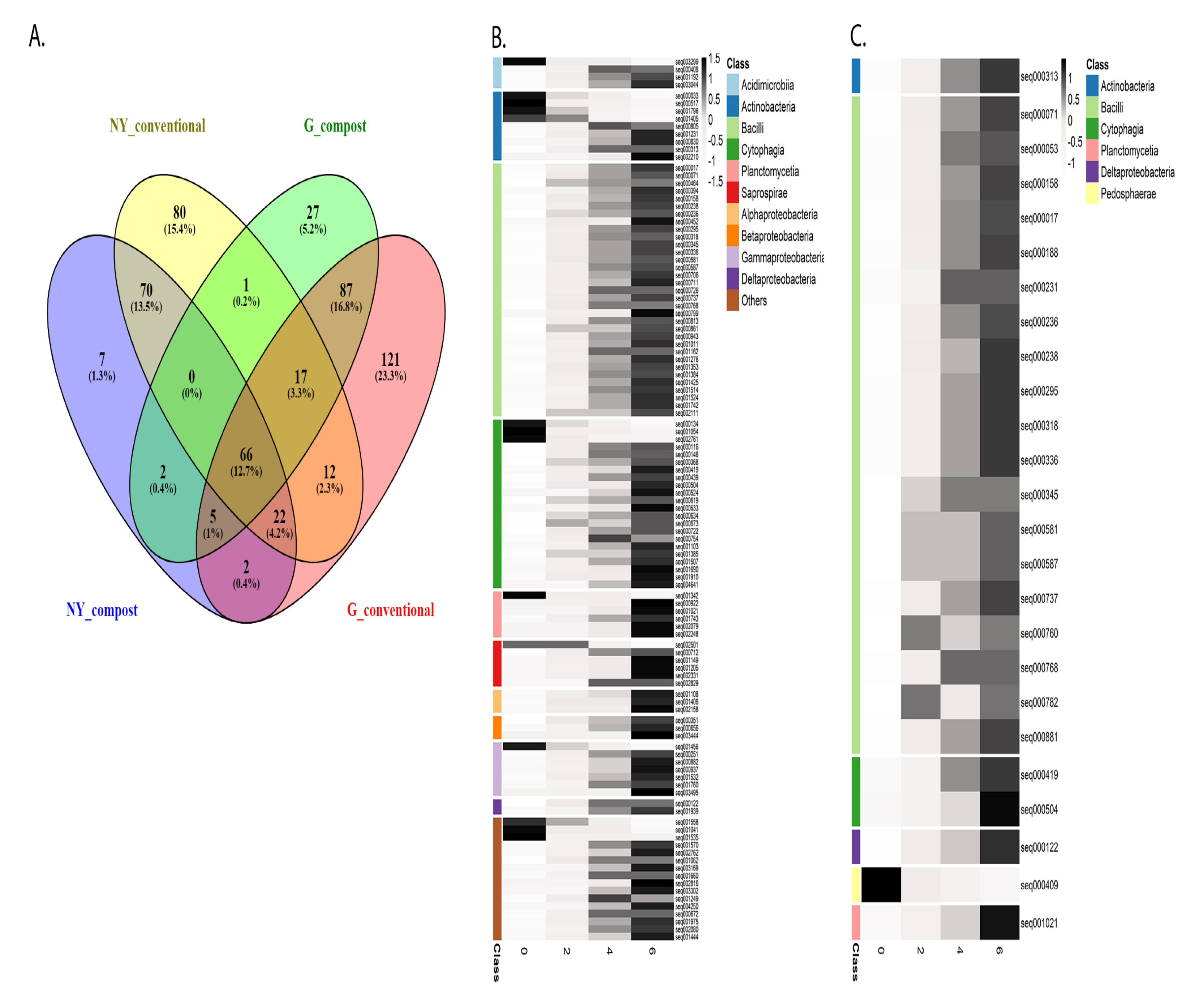
Core and indicator OTUs for compost-amended soils. (A) Venn diagram illustrating core microbiome (OTU level) of compost-amended and conventionally fertilized soils at G and NY sites. Heat map for the indicator species (significantly enriched at *P* < 0.05, OTU level) detected and shared by MaAsLin2 analysis and ANCOM (the OTU list is presented in **Table S8**) in G (B) and NY (C) soils. The heat map presents average OTU abundance at all time points for the four treatments (0, 2, 4, 6), normalized per row (Z-score). The color bars represent class-level annotation of the indicator OTUs.

To further characterize compost indicator OTUs, we analyzed the OTUs that were differentially abundant between compost and conventional soils by two methods that rely on different hypothesis tests [44]: (i) we compared all compost treatments (2, 4, 6) to conventional soil treatments (0) using MaAsLin2, and identified 1173 unique OTUs in G and 153 unique OTUs in NY; (ii) significantly differentially abundant OTUs in any treatment (0, 2, 4 or 6) were computed using ANCOM, identifying 136 OTUs in G and 36 in NY (**Table S8 A** and **B**, respectively). Finally, the lists of significant OTUs resulting from both MaAsLin2 and ANCOM analyses were compared and 111 class level-annotated OTUs in G and 25 in NY were identified by both analyses, most of which increased as a result of compost amendment. At both sites, most of these OTUs belonged to the Bacilli or Cytophagia, as well as the Planctomycetia, Deltaproteobacteria and Actinomycetota. Some of the annotated OTUs were site-specific: Pedosphaerae (higher relative abundance in the conventional treatment) in NY (**Fig. 6 C**) and Acidimicrobiia, Saprospirae, Alphaproteobacteria, Betaproteobacteria and Gammaproteobacteria (most of which increased in relative abundance in compost treatments) in G (**Fig. 6 B**). These OTUs thus represent the effect of compost on the microbiome of amended soils.

## 4. Discussion

Intensive agricultural practices require external inputs to meet crop nutrient demands. Compost is one of the most common soil amendments, and is widely used on a global scale as an alternative to mineral fertilization. Numerous studies have described the beneficial effects of organic fertilization on soil-and microbe-related parameters, such as increases in microbial biomass and total organic carbon. Nevertheless, detailed information on the immediate and prolonged effects of compost vs. conventional fertilization on the dynamics and ecology of soil microbial communities is still lacking, especially under field conditions. In this study, performed at two agricultural sites with different climates, soil types and agricultural practices, after application of different composts at each site, we characterized both the immediate and prolonged effects of conventional, chemical fertilization and three levels of compost amendment on soil chemical parameters and microbial community structure.

### 4.1. Single annual compost amendments significantly alter soil microbial communities relative to conventionally fertilized soil communities

Annual sampling between 2014 and 2016 showed that even a single compost amendment can affect microbial community structure. Similarly, previous studies have shown that a single organic soil fertilization, alone or combined with mineral fertilization, may affect microbial soil community composition [7,45,46]. In our study, compost amendment at G (soil with extremely low organic content) significantly altered the microbial community structure of agricultural soils in the first, second and third years of the experiment, in a dose-dependent manner (**Fig. 2 A**, **Table S3 E**). In NY (soil with low organic content), a significant effect was observed only in the third year of treatment, and only between the lower (treatment 2 or 4) and highest (treatment 6) compost-amended soils (**Fig. 2 B**, **Table S3 F**). In 2013, harvested vetch (*Vicia sativa*) was added to NY fields as green manure, which may have reduced the impact of the compost amendment in 2014. The compost-dependent shift in microbial community was evident in G and NY, even 8 and 10 months (respectively) after compost application (Y3 in **Fig. 2 A** and **B**). Such a prolonged effect on microbial community structure (vs. mineral fertilization or untreated soil) has been previously found after a single [47] or repeated [2,16,18,48] compost applications.

The shifts in microbial community structure associated with compost amendment in G were more closely examined; in compost-amended soils, taxa known to prefer high organic matter (e.g., Cytophagales and Myxococcales), and those described as copiotrophs and organic matter-decomposing bacteria (Bacteroidota and Bacillota) increased in relative abundance compared to the conventionally fertilized plots (**Fig. S2**). The relative abundance of Bacteroidota and Bacillota has been shown to be high during composting processes [26], and this may explain their elevated levels in the compost-amended soils. Conversely, OTUs annotated as Rhizobiales were more abundant in conventional-treatment soils (**Table S5**). G and NY soils have low organic content and originate from sites with Mediterranean or semi-arid weather conditions, with high temperatures. Interestingly, both sites have a high relative abundance of Thaumarchaeota (12.8% in NY, 8.1% in G). This phylum, which contains some ammonia-oxidizing archaea, is usually more abundant in low-fertility soils and lower in soils with higher availability of inorganic N [49–51].

### 4.2. Compost effects on soil chemistry and microbial community are dose-dependent and accumulate over time

Compost elevated many of the measured chemical parameters, mainly in G soil but also in NY soil. This is in agreement with previous studies that showed that organic fertilization increases TN, available P, TOC and pH compared to non-fertilized or mineral-fertilized soil [2,10,52,53]. This compost effect on soil chemical parameters was dose-dependent, and was most strongly observable in soils from G. For example, a significant effect of compost on the microbial community structure of G soils was visible at our first sampling point (Y1). However, this effect was stronger in 2015–2016 (Y2 and Y3 in G) or 2016 (Y3 in NY), as assessed by Bray–Curtis distances (**Fig. 2**). This enhancement of the compost effect likely reflects a cumulative effect after repeated compost amendment over several years. Alternatively, it may be affected by environmental factors (such as weather differences between Y1, Y2 and Y3; **Fig. S1**), or technical variability with different sequencing batches. When environmental and technical factors were eliminated by comparing each compost treatment to its respective conventional control (with the same environmental and technical settings), a significant difference in Y1 at the G site between treatment 2 and 6 was evident as well. Despite differences in response to amendment between the years, we observed a compost-related gradient in community composition in each of the sampling years at both sites. Such a dose-dependent change was previously observed in controlled laboratory experiments, where higher levels of compost amendment significantly affected microbial community composition (based on phospholipid fatty acid analysis or denaturing gradient gel electrophoresis profiles), while a lower dose did not [45,47]. Moreover, repeated compost application has been found to affect community structure more than a single application [3,11]. Our study demonstrates that this cumulative, dose-dependent change is also dynamic, with the community shifting over time.

### 4.3. Amendment with compost has an immediate and time-dependent effect on the soil microbiome

To follow the temporal dynamics and understand the immediate and delayed responses of the soil microbiome to compost amendment, we analyzed the microbiome dynamics at high (weekly to monthly) temporal resolution from November 2015 through 2016. This high-resolution analysis revealed significant differences in microbiome composition between conventionally fertilized (treatment 0) and compost-amended (treatments 2, 4, 6) soils, as well as between high (6) and low (2) compost doses, at both sites (**Fig. 3 A** and **B**). The observed community dissimilarity after compost application was in accordance with a previous study which found that organic and mineral fertilization explained, respectively, 15.3 and 14.4% of the dissimilarities in community composition [2]. Additionally, Saison et al. [47] showed that compost can affect microbial composition at both immediate and later (6 months) time points, in a dose-dependent manner [47].

In our study, compost application immediately shifted the soil microbial community composition compared to conventionally fertilized soils at both examined sites. This shift was reduced 4 weeks post-compost amendment and onward. In G (but not in NY), a compost dose-dependent change was also observed compared to the microbial community of conventionally fertilized soil (**Fig. 4 A** and **B**).

Compost had a different impact in each of the sites. The effect of compost dose on community composition in NY soils was milder than that in G soils, probably due to environmental parameters such as soil type and chemical properties (**Table S2**), heavy rain events (**Fig. S1**), differences between compost composition at each site as well as field management, including tillage practices, which are known to greatly impact the soil microbiome [21] (**Fig. 1**). One of these differentiating parameters is the organic carbon content, which was higher in NY soils than G soils (**Table S2**).

Our results suggest that dry, sandy soils with extremely low organic content, under semi-arid climate (as in the G field), are more susceptible to microbiome shifts resulting from agricultural management, but this finding requires further validation. Furthermore, our results suggest that frequent sampling is necessary to reliably describe the microbiome response to a disturbance such as compost amendment, as the effects of such amendments can be influenced by soil type and climate. In our case, weekly resolution was adequate for tracking immediate changes in the first month after compost amendment. To examine the delayed effect of compost, sampling between weeks 15 and 20 following compost amendment was sufficient, as the community started to stabilize at this stage.

### 4.4. Higher abundance of taxa putatively involved in organic decomposition in compost-amended soil

Organic fertilization increases soil metabolic activity and microbial functional diversity [18,54], and similar changes are likely to result from compost-mediated shifts in community structure. Our results showed that indeed, compost amendment induces taxonomic changes from the phylum to the OTU levels. At both sites, most of the OTUs that were more abundant in the compost-amended soils belonged to the phyla Bacteroidota, Pseudomonadota and Bacillota (**Fig. 4**, **Fig. S3**). Members of these bacterial groups are often involved in the degradation of complex organic compounds [9,11,55,56] and are highly abundant during composting [26]. Pseudomonadota, Bacteroidota and Bacillota are known as hydrolytic, fermentative bacteria [57,58], and have been found to be relatively more abundant in manure-fertilized soils [2,9,16,53,59]. Some of these taxonomic groups have high relative abundance in nutrient-rich environments, and Bacteroidota and Pseudomonadota have also been generally described as copiotrophic [11,55].

In NY, OTUs belonging to the phyla Actinomycetota, Chloroflexi and Crenarchaeota had higher relative abundance in conventionally fertilized soils (**Fig. S3**, **Fig. 4 D**). These results are in accordance with previous studies showing a significantly higher proportion of Actinomycetota and Chloroflexi in mineral-compared to organic-fertilized soils [2,9,16].

### 4.5. Compost-amendment indicator OTUs

Our heat map analysis showed differences in OTU composition between treatments immediately after compost amendment. Nevertheless, the relative abundance of some OTUs shifted significantly only a few weeks after compost application, thus demonstrating that compost has an additional, prolonged effect on the soil microbiome, with some OTUs being late responders. This dynamic likely reflects the balance between the fast-growing copiotrophic bacteria, which proliferate in response to the supply of readily available substrates, and the slower-growing bacteria that replace them later and utilize more recalcitrant compounds.

For the detection of compost-amendment indicator species, we identified OTUs that differ significantly in relative abundance between the compost-amended and conventionally fertilized soils. Since many tools used to detect differential abundance show variable performance between datasets, we used both MaAsLin2 and the compositionally aware method ANCOM [44]. Because ANCOM is a more conservative approach, it identifies fewer features than MaAsLin2. Most of the significantly altered OTUs were more abundant in the compost-amended soils, especially in soils amended with the highest compost levels. At both sites, most of these OTUs belonged to the class Bacilli as well as the Actinobacteria, which are known to be associated with composting processes or enriched in organically fertilized soils [48,58], and the class Cytophagia. In contrast, the relative abundance of Pedosphaerae decreased in the compost-amended soil in NY. Reduction in this class has been previously shown in soil following high-concentration N-based fertilization [60].

Core microbiome analysis identified a larger core microbiome for conventionally fertilized vs. compost-amended soils (157 vs. 30 OTUs in G, and 110 vs. 16 OTUs in NY). The smaller microbiome core size is likely the result of a more dynamic microbiome in soils with compost amendment, as also reflected in **Fig. 4 C** and **D**.

Two OTUs from the compost core were shared between the G and NY sites. Both were annotated as members of the Sinobacteraceae. This bacterial family was originally identified upon isolation of a representative member from a polluted farmland sample, and was described as a chemo-organotroph [61]. One of the two OTUs was further classified as a member of the genus *Steroidobacter*, reported to be among the most abundant genera in composting processes of dairy manure with plant additives, textiles or distilled grain waste [58,62,63].

### 4.6. Effect of compost amendment on alpha diversity of microbial communities

Some studies on long-term (21–33 years) compost application or organic fertilization have reported an increase in microbial diversity in bulk soil [18,52,64] and the rhizosphere [65], whereas others did not find fertilization-related differences in diversity [48]. Previous reports of long-and short-term compost-amendment effects on soil microbial diversity seem to be inconclusive. In some studies, bacterial diversity and richness indices were significantly higher in soils amended with organic fertilizer compared to mineral NPK-fertilized soils [2,16,18,52,64,66], whereas in others, the opposite trend was reported [10]. We could not detect a significant increase in microbial diversity in the annual sampling over 3 years (not shown), either because the sampling was done at least 8–10 months after the compost application, or due to different environments and resolution of the methodology.

However, sampling at higher temporal resolution in a single year (2016, Y3) revealed a significant compost-related effect on diversity, despite major differences in compost and soil characteristics, as well as crops and weather events between the two sites (**Tables S1** and **S2**, **Fig. S1**). The Shannon diversity index of compost-amended soils was higher throughout 2016 relative to that of conventionally fertilized soils at both sites. In NY, there was a clear effect of compost dose on the diversity, with higher compost levels leading to higher diversity (**Fig. 5 B**), whereas in G soil, diversity was similar and high at the three compost levels compared with the non-compost control (**Fig. 5 A**). NY soils were strongly affected by the compost-amendment dose compared to G soils, possibly due to the lower basal Shannon diversity in the former.

Compost amendment may increase community diversity through the addition of external microorganisms originating from the compost, or through promotion of the decomposition of complex organic matter by existing copiotrophic bacteria, the relative abundance of which is higher immediately after compost amendment. The significant difference between treatment 0 and treatments 2, 4 and 6 reflects the immediate (in the first month) fluctuations following compost amendment, as well as milder fluctuations later on, when more recalcitrant compounds are being degraded by different groups of organisms.

### 4.7. Compost-mediated site-specific differences in the microbiome

We examined microbial dynamics in two environmentally different agricultural fields that differ not only in their environmental setting (soil type, weather, rainfall) but were also amended with different commercial composts and differed in their field management (different crop rotations in G and NY and green manure applied only in NY). At all examined taxonomic levels, compost-related changes in soil microbiome composition were more significant in G than in NY. In NY, fewer OTUs were significantly enriched or depleted as a result of compost amendment. G soil had initially lower TOC and TN contents, and NO_3_, NH_4_, Cl and EC levels (**Table S2**) relative to soils of the NY site. The poor sandy soil, along with the semi-arid climate in G, likely renders this agricultural soil more susceptible to external perturbations. However, the Mediterranean site of NY, which has higher TOC and TN levels to begin with, might require a higher dose of compost to disturb its soil microbiome. NY fields were amended with harvested vetch (*Vicia sativa*) as green manure in 2013, which might have minimized the effects of the compost amendment in 2014. Previous studies have similarly demonstrated dose-dependent effects of compost amendment on rhizosphere and root microbial community structure and on root colonization [45]. Other studies have shown that the site, as a variable, can have a stronger influence on the microbiome than agricultural management [65]. Local environmental conditions may also affect the soil microbiome’s stability and responses to perturbations. For example, in our study, heavy rains in the winters of 2015 and 2016 (**Fig. S1**) may also have contributed to the reduction of compost-related effects on NY’s microbial community.

A single application of mineral fertilizer does not necessarily induce long-term changes in microbial communities [24,67]. On the other hand, continuous fertilization practice, organic or mineral, has been shown to alter soil properties and microbial community composition [16,68,69]. Continuous fertilization, i.e., more than a single fertilization event, markedly disturbs the ecological system, and may result in a new ecological steady state for the soil microbiome [2,69]. The differences between the examined sites in our study demonstrate that these long-lasting effects are site-dependent, but agricultural management and environmental variables contribute as well.

## 5. Conclusions

Results indicated that compost significantly affects soil microbiome structure immediately after amendment. This shift, which correlated to the initial applied compost dose, decayed over time. Still, some changes in the microbial community structure and ecological parameters (i.e., alpha and beta diversity) persisted for months following compost application. These changes in soil microbiome occurred in parallel to changes in soil chemical parameters. Fertilization practices were found to be more influential in shaping the soil microbiome than other site-specific or environmental factors, promoting the enrichment of microorganisms specific to compost amendment, including copiotrophic and organic matter-degrading bacteria.

## Supplementary material

Supplementary material accompanies the paper.

## Acknowledgments

This work was supported by grant no. 20-17-0004 from the Chief Scientist of the Ministry of Agriculture and Rural Development, Israel. We thank Dafna Lavi for her assistance.

## Supplementary material

**Fig. S1.** Precipitation at the sampling sites during 2014–2016. Rainfall measurements (mm) in (A) Gilat (G) and (B) Newe Ya’ar (NY) during the experimental period.

**Fig. S2.** Compost-mediated changes in relative abundance of OTU-level taxa in yearly resolution sampling. Heat map of treatment-specific OTUs calculated throughout the 3 years of sampling. All presented OTUs were significantly (Kruskal–Wallis test, FDR corrected *P* < 0.05) enriched or depleted at 0 vs. any compost treatment (2, 4 or 6). The heat map presents average OTU abundance on each sampling date for the 3 years of the study, normalized per row. The color bars correspond to phylum-level classification of the different OTUs.

**Fig. S3.** Temporal dynamics of dominant phyla representing significantly enriched or depleted OTUs in 2016, in response to compost dose. Average relative abundance of phylum-level taxa that represent all OTUs which were significantly (Kruskal–Wallis test, FDR corrected *P* < 0.05) enriched or depleted in conventional (0) vs. compost treatments (and plotted in a heat map in Fig. 4 C and D) in (G) (A) and (NY) (B) is presented. Conventional-practice (0) samples are marked in violet, and compost-amendment treatments are marked in light blue (2), blue (4), and dark blue (6).

**Fig. S4.** Comparison of core microbiomes in compost-amended vs. conventionally fertilized soils. The Venn diagram depicts the unique and shared OTUs among the core microbiomes of compost-amended vs. conventionally fertilized samples in G (A) and NY (B).

**Table S1.** Chemical and physical characteristics of compost applied during the study. Various chemical and physical properties of the compost applied in Newe Ya’ar (NY) (left) and Gilat (G) (right), as measured in 2013, and in January and December 2015.

**Table S2.** The effects of compost load on G and NY soil properties during 2014–2016. Chemical soil parameters were measured between 2014 and 2016 and analyzed by Tukey HSD test. Letters indicate statistical difference between compost treatments. Asterisks indicate measurements done in 2014–2015.

**Table S3.** Pairwise comparison between *P*-values based on year or compost parameters in the microbial community analysis during 2014–2016. Comparison of the variation between years (2014, 2015, 2016) in G (A) and NY (B) analyzed by ANOSIM. Comparison of compost treatments in all samples during 2014–2016 in G (C) and NY (D) was analyzed by ANOSIM. Dissimilarity distance comparison between treatment/year pairs in G (E) and NY (F) in each year, analyzed by PERMANOVA. Significant *P*-values are marked in red.

**Table S4.** PERMANOVA results comparing microbial communities (nMDS ordination-based distances) between all treatments in each year. Pairwise comparison using PERMANOVA to all treatment pairs (0, 2, 4, 6 indicates soils with 0, 20, 40 and 60 m^3^/ha compost amendment) in 2014 (Y1), 2015 (Y2) and 2016 (Y3) is presented after 999 permutations as shown in the nMDS ordination in Fig. 2 C and D. Significant *P*-values are marked in red.

**Table S5.** Relative abundance comparison for microbial orders between different doses of compost amendment vs. conventional practice. The relative abundance of major orders was compared between treatments in each year (2014–2016). Significant (Wilcoxon test) differential abundance between treatments is marked in red.

**Table S6.** PERMANOVA results for all treatments in G and NY during 2016. Pairwise comparison using PERMANOVA of treatment pairs (0, 2, 4, 6 indicates 0, 20, 40 and 60 m^3^/ha compost amendment) throughout all 2016 sampling time points together. Significant *P*-values are marked in red.

**Table S7.** Compost-amended and conventionally fertilized soil core microbiome features in NY and G. Shown is a list of the identified core OTUs for compost treatments in G, NY or both, as well as core OTUs that are shared between conventional treatments in NY and G.

**Table S8.** Significant OTUs identified by MaAsLin2 and ANCOM. List of significant OTUs that were identified by ANCOM, MaAsLin2, or by both methods, are presented for G (A) and NY (B) sites.

