## Supplementary figures for "Short- and long-term effects of continuous compost amendment on soil microbiome community"

### Supplementary Figures and Tables

Cohen et al 2023

Fig. S1

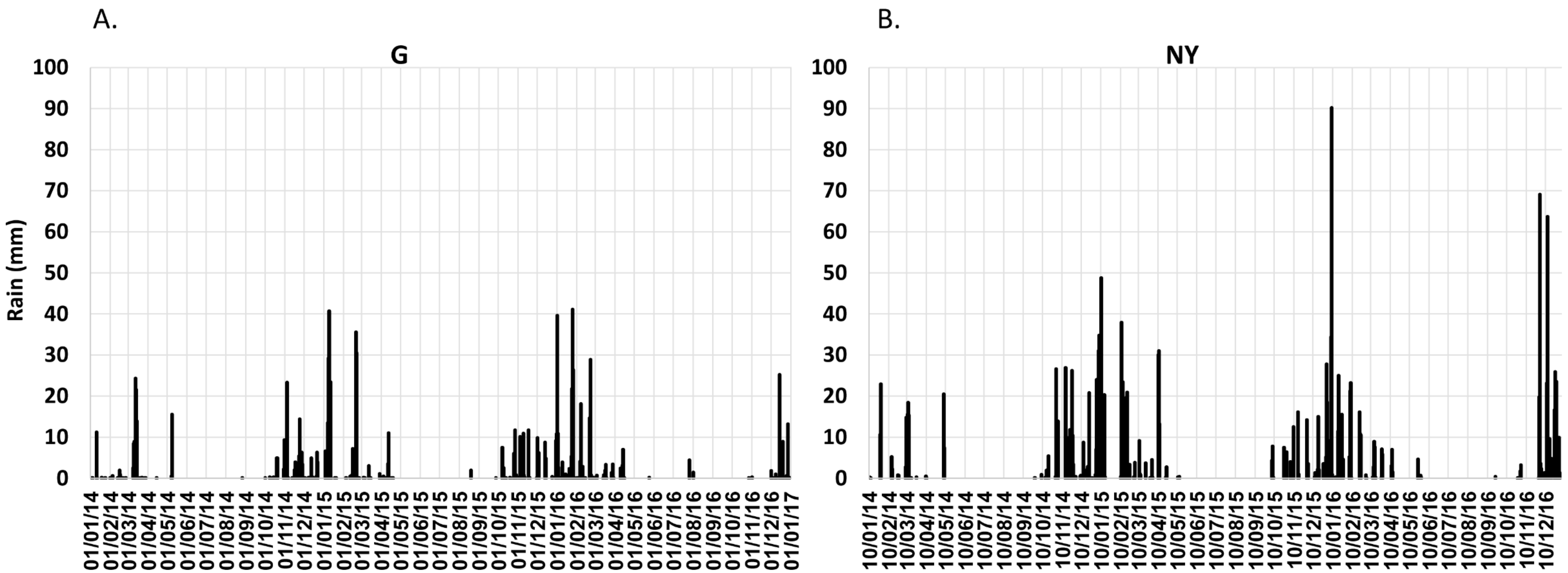
A.
G
B.
NY
Rain (mm)
100
90
80
70
60
50
40
30
20
10
0
01/01/14
01/02/14
01/03/14
01/04/14
01/05/14
01/06/14
01/07/14
01/08/14
01/09/14
01/10/14
01/11/14
01/12/14
01/01/15
01/02/15
01/03/15
01/04/15
01/05/15
01/06/15
01/07/15
01/08/15
01/09/15
01/10/15
01/11/15
01/12/15
01/01/16
01/02/16
01/03/16
01/04/16
01/05/16
01/06/16
01/07/16
01/08/16
01/09/16
01/10/16
01/11/16
01/12/16
01/01/17
10/01/14
10/02/14
10/03/14
10/04/14
10/05/14
10/06/14
10/07/14
10/08/14
10/09/14
10/10/14
10/11/14
10/12/14
10/01/15
10/02/15
10/03/15
10/04/15
10/05/15
10/06/15
10/07/15
10/08/15
10/09/15
10/10/15
10/11/15
10/12/15
10/01/16
10/02/16
10/03/16
10/04/16
10/05/16
10/06/16
10/07/16
10/08/16
10/09/16
10/10/16
10/11/16
10/12/16

Fig. S2

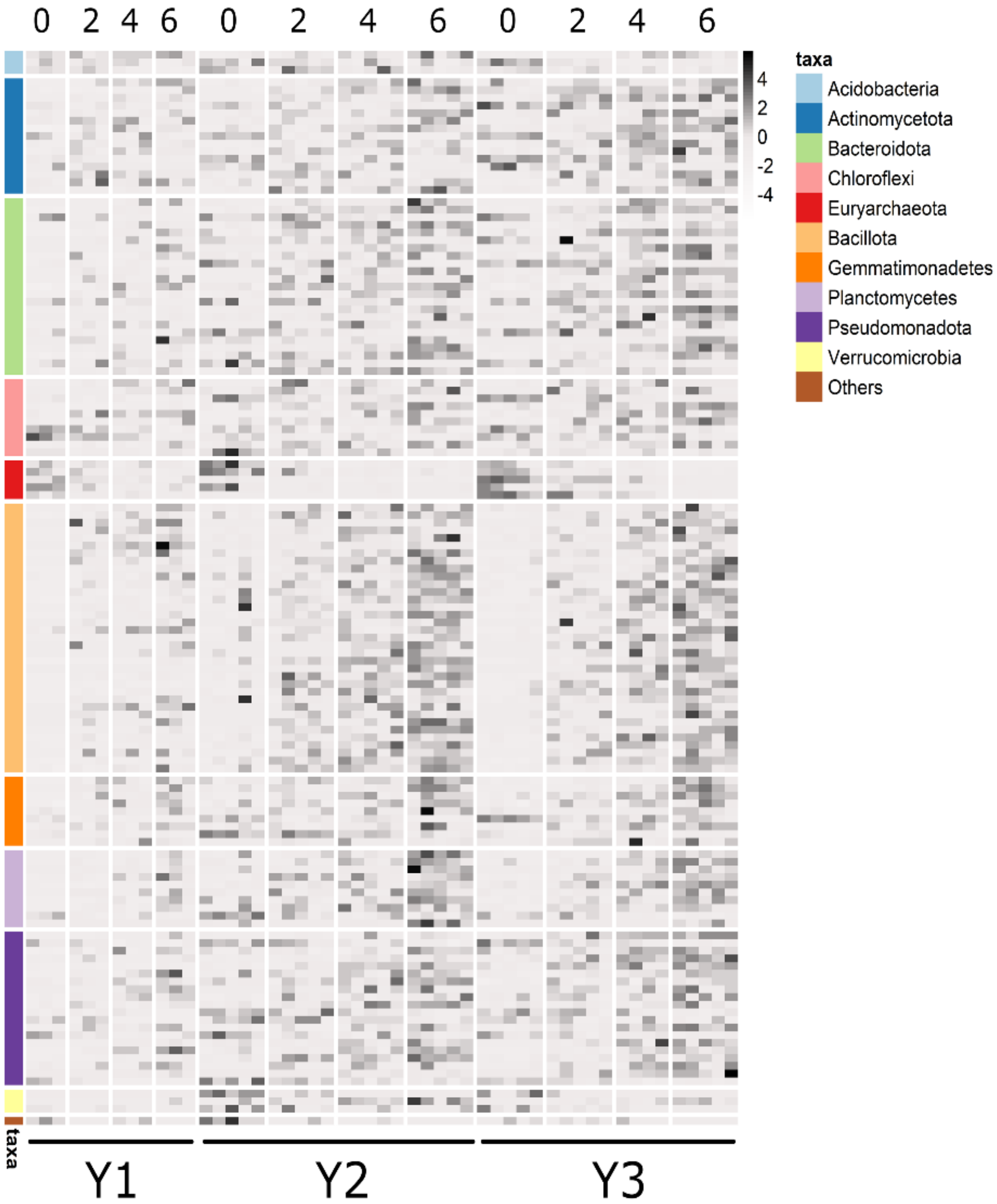
0
2
4
6
0
2
4
6
0
2
4
6
4
2
0
-2
-4
taxa
Acidobacteria
Actinomycetota
Bacteroidota
Chloroflexi
Euryarchaeota
Bacillota
Gemmatimonadetes
Planctomycetes
Pseudomonadota
Verrucomicrobia
Others
taxa
Y1
Y2
Y3

Fig. S3

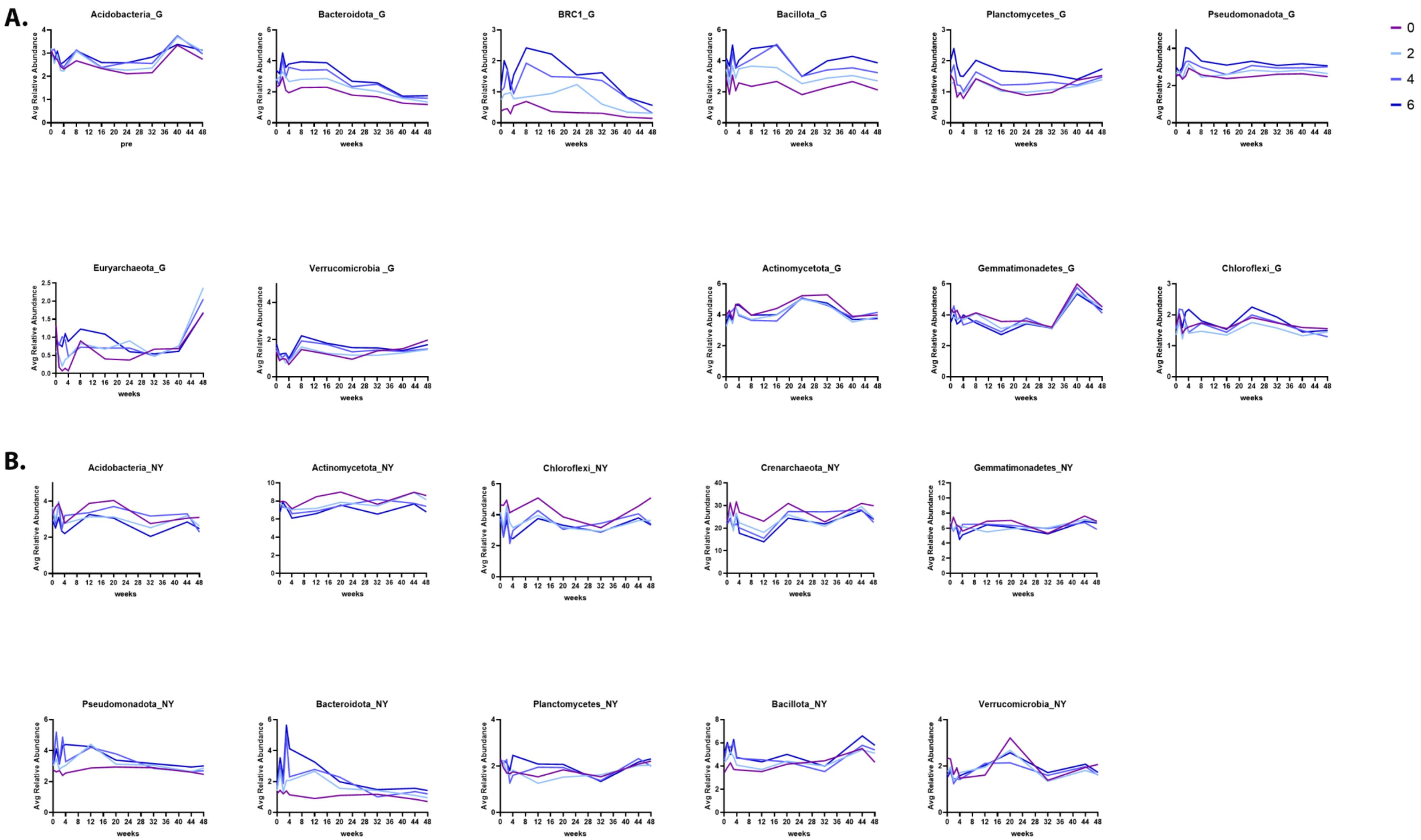
A.
Acidobacteria_G
Bacteroidota_G
BRC1_G
Bacillota_G
Planctomycetes_G
Pseudomonadota_G
Euryarchaeota_G
Verrucomicrobia _G
Actinomycetota_G
Gemmatimonadetes_G
Chloroflexi_G
B.
Acidobacteria_NY
Actinomycetota_NY
Chloroflexi_NY
Crenarchaeota_NY
Gemmatimonadetes_NY
Pseudomonadota_NY
Bacteroidota_NY
Planctomycetes_NY
Bacillota_NY
Verrucomicrobia_NY
Avg Relative Abundance
weeks
pre
0
2
4
6

Fig. S4

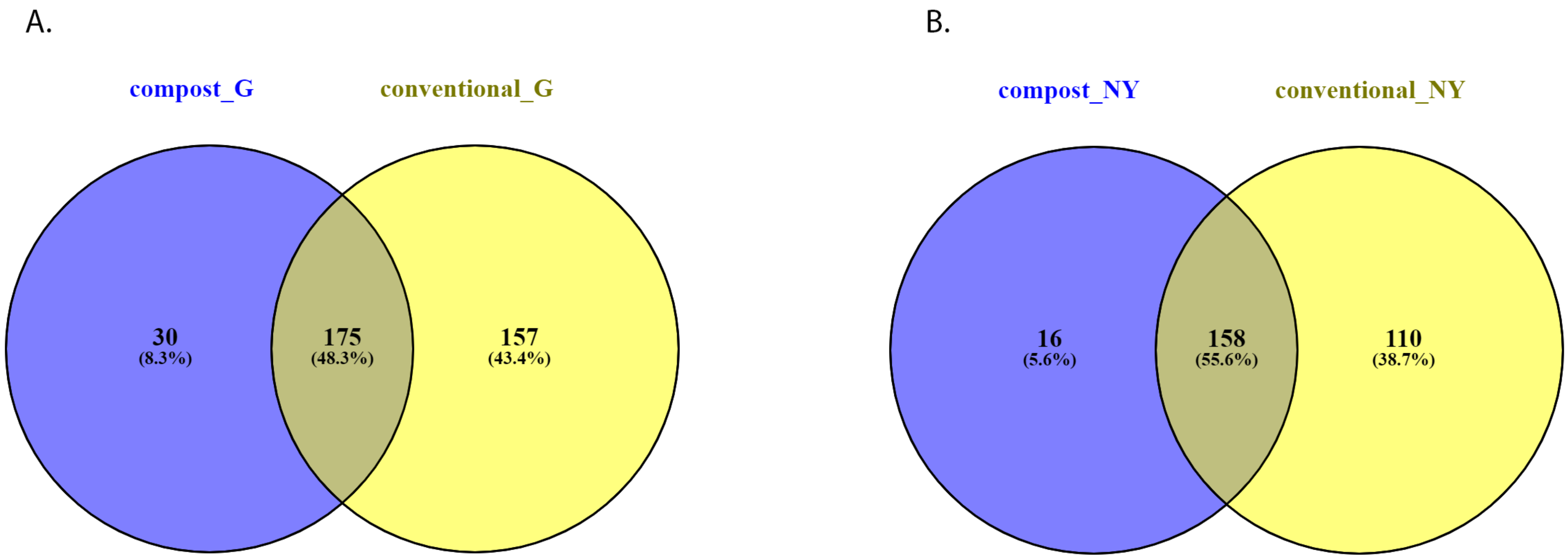
A.
compost_G
conventional_G
30
(8.3%)
175
(48.3%)
157
(43.4%)
B.
compost_NY
conventional_NY
16
(5.6%)
158
(55.6%)
110
(38.7%)

Table S1- Chemical and physical characteristics of compost applied during the study

| | | G | | | NY | |
|---|---|---|---|---|---|---|
| Variable | Units | 2013 | 2015 (January) | 2015 (December) | 2014 | 2015 |
| Organic C | % w/w | 16.6 | 31.7 | 14.4 | 25.9 | 30.1 |
| Total N | % w/w | 1.3 | 2.0 | 1.0 | 1.7 | 2.0 |
| C/N | | 12.6 | 16.0 | 14.2 | 12.9 | 15.6 |
| EC | dS m-1 | 8.3 | 8.9 | 5.1 | 9.7 | 8.1 |
| N-NO3- | mg kg-1 | 319.0 | 18.0 | 634.0 | 34.0 | 94.0 |
| N-NH4+ | mg kg-1 | 41.0 | 1261.5 | 165.5 | 168.0 | 218.0 |
| P | g kg-1 | 0.3 | 2.8 | 0.4 | 0.7 | 0.3 |
| K | g kg-1 | 11.8 | 7.9 | 8.5 | 17.6 | 15.5 |
| N-NO3-/N-NH4+ | | 7.8 | 0.0 | 3.8 | 0.2 | 0.4 |
| Bulk density | $kg/m^3$ | 667.0 | 338.0 | 636.0 | 488.0 | 383.0 |
| Ash | % w/w | 71.0 | 45.0 | 75.0 | 54.0 | 48.0 |
| pH | | 8.8 | 6.7 | NT | 7.2 | NT |

NT=not tested

The chemical tests for all composts were done at the "Newe Ya'ar field service laboratory" according to standard methods

Table S2. The effects of compost load on Gilat and Newe Ya'ar soil properties during 2014-2016.

| Variable | TOC | TN | NH4 | NO3 | P | K | Na | Cl | EC | pH |
|---|---|---|---|---|---|---|---|---|---|---|
| | mg g$^{-1}$ | | mg kg$^{1-}$ | | | | | | dS m$^{-1}$ | |
| | Gilat | | | | | | | | | |
| Compost Dose | | | | | | | | | | |
| 0 | 7.19 c | 0.719 d | 9.08 | 11.0 b | 11.1 d | 61.3 c | 60.7 c | 48.1 | 0.175 b | 7.28 |
| 2 | 9.50 b | 0.975 c | 8.83 | 11.9 b | 54.9 c | 126.0 b | 88.7 b | 60 | 0.187 b | 7.37 |
| 4 | 10.23 b | 1.087 b | 9.2 | 18.7 ab | 90.2 b | 163.7 ab | 112.0 ab | 56.6 | 0.224 ab | 7.24 |
| 6 | 12.26 a | 1.297 a | 9.68 | 22.5 a | 123.9 a | 190.9 a | 132.0 a | 66.7 | 0.251 a | 7.32 |
| | Newe Ya'ar | | | | | | | | | |
| Compost Dose | | | | | | | | | | |
| 0 | 18.39 | 1.69 b | 13.49 | 18.3 | 15.2 a | 28.2 b | | 170.2 | 0.237 | 7.21 |
| 2 | 19.51 | 1.85 ab | 14.3 | 20.2 | 41.0 b | 48.0 a | | 172.2 | 0.265 | 7.11 |
| 4 | 20.99 | 1.98 ab | 14.48 | 19.7 | 46.8 b | 54.4 a | | 146.4 | 0.269 | 7.2 |
| 6 | 20.27* | 2.02* a | 14.31 | 23.1 | 67.8 c | 46.9 a | | 139.4 | 0.256 | 7.19 |

*Averge data from years 2014-2015

Table S3 -ANOSIM analysis to dissimilarity distance treatment/year pairs in Gilat and NY 2014-2016

| | Site | Treatment1 | Treatment2 | p-value | q-value |
|---|---|---|---|---|---|
| A | G | Y1 | Y2 | 0.001 | 0.001 |
| | | Y1 | Y3 | 0.001 | 0.001 |
| | | Y2 | Y3 | 0.001 | 0.001 |
| B | NY | Y1 | Y2 | 0.001 | 0.001 |
| | | Y1 | Y3 | 0.001 | 0.001 |
| | | Y2 | Y3 | 0.001 | 0.001 |

| | Site | Treatment1 | Treatment2 | p-value | q-value |
|---|---|---|---|---|---|
| C | G | 0 | 2 | 0.131 | 0.1965 |
| | | 0 | 4 | 0.013 | 0.039 |
| | | 0 | 6 | 0.001 | 0.006 |
| | | 2 | 4 | 0.536 | 0.6432 |
| | | 2 | 6 | 0.051 | 0.102 |
| | | 4 | 6 | 0.649 | 0.649 |
| D | NY | 0 | 2 | 0.525 | 0.7875 |
| | | 0 | 4 | 0.123 | 0.369 |
| | | 0 | 6 | 0.105 | 0.369 |
| | | 2 | 4 | 0.837 | 0.972 |
| | | 2 | 6 | 0.36 | 0.72 |
| | | 4 | 6 | 0.972 | 0.972 |

Kruskal Wallis analysis to dissimilarity distance treatment pairs per year in Gilat and NY 2014-2016

E

| G-Year | Treatment1 | Treatment2 | p-Value |
|---|---|---|---|
| Y1 | 6 | 2 | 0.017 |
| Y1 | 4 | 2 | 0.064 |
| Y1 | 6 | 4 | 0.289 |
| Y2 | 6 | 2 | 0.0001 |
| Y2 | 6 | 4 | 0.0008 |
| Y2 | 4 | 2 | 0.2687 |
| Y3 | 6 | 2 | 0.0001 |
| Y3 | 6 | 4 | 0.0001 |
| Y3 | 4 | 2 | 0.0001 |

F

| NY-Year | Treatment1 | Treatment2 | p-Value |
|---|---|---|---|
| Y1 | 6 | 2 | 0.331 |
| Y1 | 4 | 2 | 0.377 |
| Y1 | 6 | 4 | 0.724 |
| Y2 | 6 | 2 | 0.125 |
| Y2 | 4 | 2 | 0.213 |
| Y2 | 6 | 4 | 0.868 |
| Y3 | 6 | 2 | 0.012 |
| Y3 | 6 | 4 | 0.016 |

Table S4 Permanova analysis to NMDS for treatment pairs Gilat and NY 2014-2016

| treatment comparison | test statistic | p-value | treatments |
|---|---|---|---|
| G_Y1_0_2 | 1.157 | 0.295 | 0-2 |
| G_Y1_0_4 | 1.407 | 0.203 | 0-4 |
| G_Y1_0_6 | 1.843 | 0.106 | 0-6 |
| G_Y1_2_4 | 0.906 | 0.594 | 2-4 |
| G_Y1_2_6 | 1.091 | 0.183 | 2-6 |
| G_Y1_4-6 | 0.986 | 0.596 | 4-6 |
| G_Y2_0_2 | 1.293 | 0.106 | 0-2 |
| G_Y2_0_4 | 1.783 | 0.029 | 0-4 |
| G_Y2_0_6 | 2.924 | 0.008 | 0-6 |
| G_Y2_2_4 | 1.123 | 0.211 | 2-4 |
| G_Y2_2_6 | 1.650 | 0.008 | 2-6 |
| G_Y2_4_6 | 1.210 | 0.097 | 4-6 |
| G_Y3_0_2 | 1.714 | 0.009 | 0-2 |
| G_Y3_0_4 | 2.433 | 0.008 | 0-4 |
| G_Y3_0_6 | 3.483 | 0.01 | 0-6 |
| G_Y3_2_4 | 1.136 | 0.145 | 2-4 |
| G_Y3_2_6 | 1.924 | 0.009 | 2-6 |
| G_Y3_4_6 | 0.968 | 0.521 | 4-6 |
| **treatment comparison** | **test statistic** | **p-value** | **treatments** |
| NY_Y1_0_2 | 1.091 | 0.183 | 0-2 |
| NY_Y1_0_4 | 1.027 | 0.373 | 0-4 |
| NY_Y1_0_6 | 1.090 | 0.293 | 0-6 |
| NY_Y1_2_4 | 1.091 | 0.183 | 2-4 |
| NY_Y1_2_6 | 0.845 | 0.902 | 2-6 |
| NY_Y1_4_6 | 0.935 | 0.495 | 4-6 |
| NY_Y2_0_2 | 0.856 | 0.478 | 0-2 |
| NY_Y2_0_4 | 1.195 | 0.173 | 0-4 |
| NY_Y2_0_6 | 1.414 | 0.145 | 0-6 |
| NY_Y2_2_4 | 0.821 | 0.579 | 2-4 |
| NY_Y2_2_6 | 1.057 | 0.25 | 2-6 |
| NY_Y2_4_6 | 0.890 | 0.437 | 4-6 |
| NY_Y3_0_2 | 0.811 | 0.867 | 0-2 |
| NY_Y3_0_4 | 1.369 | 0.075 | 0-4 |
| NY_Y3_0_6 | 1.396 | 0.083 | 0-6 |
| NY_Y3_2_4 | 1.106 | 0.153 | 2-4 |
| NY_Y3_2_6 | 0.975 | 0.49 | 2-6 |
| NY_Y3_4_6 | 0.791 | 0.913 | 4-6 |

Table S5-Order level relative abundance comparison between treatments of different compost amendment doses and conventional practice

| Order | G_Y1_0 | G_Y1_2 | G_Y1_4 | G_Y1_6 | G_Y2_0 | G_Y2_2 | G_Y2_4 | G_Y2_6 | G_Y3_0 | G_Y3_2 | G_Y3_4 | G_Y3_6 |
|---|---|---|---|---|---|---|---|---|---|---|---|---|
| **Bacillales** | 0.131 | 0.152 | 0.145 | 0.158 | 0.122 | 0.131 | 0.143 | 0.140 | 0.102 | 0.110 | 0.113 | 0.129 |
| **Rhizobiales** | 0.099 | 0.093 | 0.102 | 0.103 | 0.063 | 0.066 | 0.062 | 0.067 | 0.082 | 0.081 | 0.078 | 0.075 |
| **Micrococcales** | 0.084 | 0.080 | 0.100 | 0.082 | 0.050 | 0.044 | 0.048 | 0.046 | 0.073 | 0.062 | 0.065 | 0.062 |
| **Cytophagales** | 0.022 | 0.029 | 0.034 | 0.034 | 0.052 | 0.063 | 0.061 | 0.070 | 0.037 | 0.041 | 0.047 | 0.047 |
| **Sphingobacteriales** | 0.019 | 0.027 | 0.029 | 0.024 | 0.056 | 0.062 | 0.056 | 0.059 | 0.042 | 0.046 | 0.047 | 0.044 |
| **Gemmatimonadales** | 0.047 | 0.042 | 0.037 | 0.040 | 0.037 | 0.033 | 0.032 | 0.028 | 0.034 | 0.031 | 0.029 | 0.028 |
| **Burkholderiales** | 0.040 | 0.039 | 0.065 | 0.036 | 0.024 | 0.027 | 0.026 | 0.035 | 0.032 | 0.028 | 0.030 | 0.026 |
| **Rhodospirillales** | 0.032 | 0.029 | 0.027 | 0.032 | 0.030 | 0.029 | 0.030 | 0.028 | 0.038 | 0.039 | 0.036 | 0.035 |
| **Xanthomonadales** | 0.025 | 0.027 | 0.026 | 0.026 | 0.032 | 0.039 | 0.035 | 0.032 | 0.029 | 0.030 | 0.032 | 0.030 |
| **Soil Crenarchaeotic Group(SCG)** | 0.042 | 0.040 | 0.031 | 0.032 | 0.036 | 0.035 | 0.031 | 0.028 | 0.023 | 0.028 | 0.023 | 0.020 |
| **Solirubrobacterales** | 0.036 | 0.029 | 0.027 | 0.032 | 0.024 | 0.019 | 0.021 | 0.019 | 0.035 | 0.030 | 0.032 | 0.033 |
| **Myxococcales** | 0.024 | 0.024 | 0.027 | 0.029 | 0.023 | 0.023 | 0.023 | 0.021 | 0.029 | 0.033 | 0.034 | 0.035 |
| **Sphingomonadales** | 0.025 | 0.027 | 0.028 | 0.024 | 0.022 | 0.026 | 0.022 | 0.022 | 0.028 | 0.025 | 0.026 | 0.028 |
| **Rubrobacterales** | 0.029 | 0.021 | 0.019 | 0.017 | 0.026 | 0.022 | 0.023 | 0.019 | 0.025 | 0.024 | 0.021 | 0.019 |
| **Planctomycetales** | 0.016 | 0.018 | 0.016 | 0.022 | 0.024 | 0.023 | 0.024 | 0.026 | 0.021 | 0.022 | 0.024 | 0.023 |
| **Soil Crenarchaeotic Group(SCG);uncultured crenarchaeote** | 0.024 | 0.023 | 0.017 | 0.019 | 0.023 | 0.021 | 0.021 | 0.023 | 0.016 | 0.019 | 0.018 | 0.019 |
| **others** | 0.304 | 0.298 | 0.271 | 0.290 | 0.357 | 0.340 | 0.342 | 0.337 | 0.353 | 0.350 | 0.345 | 0.347 |

Table S6- Permanova analysis for bray-curtis distances based PCoA 2016

G

| Group 1 | Group 2 | Sample size | pseudo-F | p-value | q-value |
|---|---|---|---|---|---|
| 2 | 4 | 109 | 1.84 | 0.006 | 0.006 |
| 2 | 6 | 110 | 5.35 | 0.001 | 0.0012 |
| 2 | 0 | 109 | 4.46 | 0.001 | 0.0012 |
| 4 | 6 | 109 | 2.35 | 0.001 | 0.0012 |
| 4 | 0 | 108 | 7.61 | 0.001 | 0.0012 |
| 6 | 0 | 109 | 13.48 | 0.001 | 0.0012 |

NY

| Group 1 | Group 2 | Sample size | pseudo-F | p-value | q-value |
|---|---|---|---|---|---|
| 2 | 4 | 100 | 1.25 | 0.06 | 0.072 |
| 2 | 6 | 100 | 2.34 | 0.001 | 0.0015 |
| 2 | 0 | 98 | 2.44 | 0.001 | 0.0015 |
| 4 | 6 | 100 | 1.21 | 0.099 | 0.099 |
| 4 | 0 | 98 | 3.88 | 0.001 | 0.0015 |
| 6 | 0 | 98 | 6.17 | 0.001 | 0.0015 |

### Table S7- Core features analysis

| Compost related core features in G | | |
|---|---|---|
| Plot | OUT no. | Closest annotation |
| G | seq000071 | *Bacillaceae* |
| G | seq000087 | *Sinobacteraceae* |
| G | seq000116 | *Pontibacter* |
| G | seq000119 | *Steroidobacter* |
| G | seq000122 | *Myxococcales* |
| G | seq000128 | *Streptomyces* |
| G | seq000141 | *Paenibacillus amylolyticus* |
| G | seq000158 | *Thermoactinomycetaceae* |
| G | seq000169 | *Steroidobacter* |
| G | seq000189 | *Ellin517* |
| G | seq000191 | *Planococcus* |
| G | seq000236 | *Bacillaceae* |
| G | seq000238 | *Bacillus* |
| G | seq000257 | *Bacillus* |
| G | seq000297 | *Bacillus firmus* |
| G | seq000299 | *Planctomyces* |
| G | seq000345 | *Virgibacillus* |
| G | seq000368 | *Pontibacter* |
| G | seq000391 | *Chthoniobacteraceae* |
| G | seq000430 | *Blastococcus aggregatus* |
| G | seq000439 | *Pontibacter* |
| G | seq000451 | *Ellin6075* |
| G | seq000499 | *Ellin6075* |
| G | seq000555 | *Sphingobacteriales* |
| G | seq000601 | *Geodermatophilus* |
| G | seq000659 | *Piscirickettsiaceae* |
| G | seq000729 | *Candidatus Nitrososphaera SCA1145* |
| G | seq000744 | *Azohydromonas* |
| G | seq000748 | *Devosia* |
| G | seq000767 | *Kouleothrixaceae* |

| Compost related core features in NY | | |
|---|---|---|
| NY | seq000049 | Sinorhizobium |
| NY | seq000100 | Pontibacter |
| NY | seq000196 | Rubrobacter |
| NY | seq000087 | Sinobacteraceae |
| NY | seq000259 | Candidatus Nitrososphaera |
| NY | seq000132 | Thermomonas |
| NY | seq000209 | Nitrososphaera gargensis |
| NY | seq000176 | Ellin6075 |
| NY | seq000261 | Rubrobacteraceae |
| NY | seq000402 | Gaiellaceae |
| NY | seq000119 | Steroidobacter |
| NY | seq000093 | Ellin6529 |
| NY | seq000149 | Bacillus selenatarsenatis |
| NY | seq000198 | Balneimonas |
| NY | seq000292 | Rubrobacter |
| NY | seq000104 | Streptomyces reticuliscabiei |

| Compost related core features in G and NY | | |
|---|---|---|
| NY and G | seq000087 | Sinobacteraceae |
| NY and G | seq000119 | Steroidobacter |

| Conventional related core features in G and NY | | |
|---|---|---|
| NY and G | seq000159 | Geodermatophilus |
| NY and G | seq000184 | WD2101 |
| NY and G | seq000414 | iii1-15 |
| NY and G | seq000286 | iii1-15 |
| NY and G | seq000171 | Rhizobiales |
| NY and G | seq000327 | Modestobacter |
| NY and G | seq000406 | Nitrospiraceae |
| NY and G | seq002035 | Skermanella |
| NY and G | seq000288 | Planococcaceae |
| NY and G | seq000194 | Comamonadaceae |
| NY and G | seq000202 | Gemm-1 |
| NY and G | seq000285 | Bacillales |

**Table S8A -Significant OTUs identified by MaAsLin2 and ANCOM G**

| Ancom G | MaAsLin2 G | Common elements G |
|---|---|---|
| seq003612 | seq000017 | seq000017 |
| seq000408 | seq000017 | seq000071 |
| seq000922 | seq000071 | seq000158 |
| seq000251 | seq000071 | seq000295 |
| seq002331 | seq000158 | seq000033 |
| seq004317 | seq000295 | seq000251 |
| seq000633 | seq000033 | seq000394 |
| seq000415 | seq000251 | seq000236 |
| seq000076 | seq000394 | seq000238 |
| seq000706 | seq000071 | seq000318 |
| seq001103 | seq000158 | seq000134 |
| seq000158 | seq000037 | seq000076 |
| seq000071 | seq000236 | seq000345 |
| seq001021 | seq000251 | seq000368 |
| seq002816 | seq000017 | seq000922 |
| seq000345 | seq000295 | seq001041 |
| seq000238 | seq000238 | seq000524 |
| seq001760 | seq000318 | seq000633 |
| seq000295 | seq000134 | seq001535 |
| seq000394 | seq000076 | seq001384 |
| seq000937 | seq000345 | seq001149 |
| seq002397 | seq000368 | seq000799 |
| seq000236 | seq000236 | seq000116 |
| seq002762 | seq000174 | seq001108 |
| seq002501 | seq000922 | seq000415 |
| seq000805 | seq000318 | seq001514 |
| seq001108 | seq001041 | seq001342 |
| seq000581 | seq000238 | seq000656 |
| seq002079 | seq000158 | seq001796 |
| seq002142 | seq000027 | seq001231 |
| seq000318 | seq000394 | seq001690 |
| seq000813 | seq000524 | seq000336 |
| seq000341 | seq000633 | seq001021 |
| seq000829 | seq000168 | seq000419 |
| seq001690 | seq000174 | seq000517 |
| seq003424 | seq001535 | seq000452 |
| seq001796 | seq000033 | seq000351 |
| seq000587 | seq001384 | seq000122 |
| seq001342 | seq001149 | seq000587 |
| seq000830 | seq000305 | seq000813 |
| seq000799 | seq000799 | seq001444 |
| seq000017 | seq000171 | seq001205 |
| seq000452 | seq000499 | seq001456 |
| seq001249 | seq000116 | seq001507 |
| seq000464 | seq001108 | seq000495 |
| seq001192 | seq000169 | seq000672 |
| seq004424 | seq000415 | seq000439 |
| seq001162 | seq000193 | seq000464 |
| seq000711 | seq001514 | seq001054 |
| seq000726 | seq001377 | seq000722 |
| seq001149 | seq000037 | seq000146 |
| seq001939 | seq001342 | seq001405 |
| seq009299 | seq000656 | seq000937 |
| seq001507 | seq000013 | seq001910 |
| seq000768 | seq000137 | seq001353 |
| seq003444 | seq000236 | seq000726 |
| seq000134 | seq001796 | seq002501 |
| seq004188 | seq000009 | seq001743 |
| seq001041 | seq001231 | seq001558 |
| seq001570 | seq001690 | seq001975 |

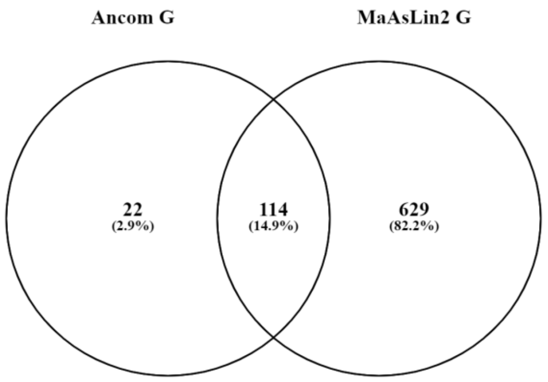

| | | |
|---|---|---|
| seq000943 | seq001342 | seq000634 |
| seq000789 | seq000336 | seq000861 |
| seq007362 | seq000104 | seq001570 |
| seq000116 | seq000076 | seq000711 |
| seq000673 | seq001535 | seq000619 |
| seq002080 | seq001342 | seq001408 |
| seq001929 | seq000368 | seq002761 |
| seq002761 | seq000269 | seq003299 |
| seq000634 | seq000169 | seq000712 |
| seq001975 | seq001384 | seq001760 |
| seq000524 | seq000541 | seq001524 |
| seq001353 | seq001535 | seq001425 |
| seq000336 | seq000208 | seq002248 |
| seq002248 | seq000116 | seq002762 |
| seq001054 | seq001021 | seq002331 |
| seq001276 | seq000419 | seq000737 |
| seq002829 | seq000600 | seq000673 |
| seq001205 | seq001435 | seq001385 |
| seq001742 | seq000517 | seq000754 |
| seq001535 | seq000462 | seq002210 |
| seq001062 | seq000452 | seq000504 |
| seq000313 | seq000043 | seq001011 |
| seq000882 | seq000264 | seq001103 |
| seq004641 | seq000594 | seq001162 |
| seq000737 | seq000351 | seq002080 |
| seq001444 | seq000020 | seq000313 |
| seq002111 | seq000122 | seq001532 |
| seq000419 | seq000192 | seq001742 |
| seq000033 | seq000587 | seq001660 |
| seq000722 | seq000988 | seq000805 |
| seq000368 | seq000813 | seq001062 |
| seq001743 | seq001377 | seq002079 |
| seq002612 | seq001391 | seq000943 |
| seq002197 | seq000597 | seq001939 |
| seq001524 | seq000081 | seq002111 |
| seq001660 | seq000073 | seq000581 |
| seq000504 | seq000062 | seq003495 |
| seq001910 | seq000295 | seq000830 |
| seq001384 | seq000027 | seq000706 |
| seq002147 | seq000415 | seq001192 |
| seq000517 | seq000122 | seq003169 |
| seq001425 | seq000584 | seq001276 |
| seq001011 | seq000076 | seq000882 |
| seq000619 | seq000302 | seq000768 |
| seq000351 | seq001444 | seq004641 |
| seq004250 | seq000104 | seq002816 |
| seq000754 | seq000016 | seq001249 |
| seq001405 | seq000368 | seq003444 |
| seq002116 | seq001205 | seq002158 |
| seq001456 | seq000188 | seq000408 |
| seq005116 | seq001456 | seq003302 |
| seq000495 | seq000066 | seq003044 |
| seq003044 | seq000664 | seq004250 |
| seq000712 | seq000318 | seq002829 |
| seq000146 | seq000269 | |
| seq000122 | seq001507 | |
| seq000672 | seq000495 | |
| seq001385 | seq000013 | |
| seq002210 | seq000672 | |
| seq001231 | seq000863 | |
| seq002158 | seq000326 | |
| seq000439 | seq001560 | |

| | |
|---|---|
| seq001514 | seq002899 |
| seq003495 | seq000439 |
| seq001558 | seq000345 |
| seq002822 | seq000143 |
| seq000656 | seq001698 |
| seq003299 | seq000977 |
| seq000861 | seq000210 |
| seq002150 | seq000653 |
| seq001409 | seq000704 |
| seq003302 | seq001041 |
| seq003169 | seq000188 |
| seq000782 | seq000464 |
| seq001532 | seq001054 |
| seq001408 | seq000115 |
| | seq000004 |
| | seq000653 |
| | seq000506 |
| | seq000722 |
| | seq000146 |
| | seq000231 |
| | seq000043 |
| | seq000009 |
| | seq001405 |
| | seq000251 |
| | seq000208 |
| | seq000509 |
| | seq000625 |
| | seq000043 |
| | seq001514 |
| | seq000146 |
| | seq000484 |
| | seq000517 |
| | seq000033 |
| | seq000937 |
| | seq001910 |
| | seq001177 |
| | seq000270 |
| | seq000508 |
| | seq000776 |
| | seq000464 |
| | seq001397 |
| | seq000672 |
| | seq001353 |
| | seq004704 |
| | seq000816 |
| | seq000238 |
| | seq000336 |
| | seq000726 |
| | seq000087 |
| | seq000305 |
| | seq000299 |
| | seq002501 |
| | seq000351 |
| | seq000122 |
| | seq000707 |
| | seq000210 |
| | seq000292 |
| | seq000767 |
| | seq000169 |
| | seq001743 |
| | seq001259 |
| | seq000499 |

seq000116
seq000138
seq001558
seq000269
seq000078
seq001975
seq001405
seq002899
seq000813
seq000611
seq000634
seq002665
seq000861
seq000134
seq001059
seq000524
seq000020
seq000877
seq004704
seq000393
seq001570
seq000430
seq004470
seq003597
seq000650
seq000711
seq000729
seq002800
seq000619
seq002743
seq000464
seq001408
seq000105
seq000462
seq001607
seq002761
seq002483
seq001153
seq000307
seq003299
seq001397
seq000676
seq000712
seq000451
seq001760
seq000722
seq000462
seq000014
seq000034
seq001248
seq002264
seq000270
seq000439
seq003431
seq000712
seq000166
seq000452
seq002554
seq000260
seq000013
seq000393
seq000003

seq001041
seq001524
seq003500
seq002761
seq002818
seq002280
seq004470
seq001677
seq001425
seq000595
seq000045
seq002576
seq000168
seq003113
seq000495
seq000611
seq000443
seq001042
seq004704
seq000421
seq000194
seq000491
seq000840
seq002787
seq001102
seq000541
seq000166
seq001925
seq001196
seq000451
seq000925
seq000909
seq000383
seq001796
seq001029
seq000780
seq001054
seq000925
seq000854
seq000665
seq002248
seq000001
seq001183
seq002762
seq000463
seq000834
seq000669
seq000364
seq000503
seq000671
seq000555
seq001261
seq001456
seq002331
seq000357
seq000099
seq000081
seq001992
seq000737
seq000673
seq002501
seq000676

seq001219
seq000414
seq000015
seq002056
seq000222
seq000171
seq001290
seq000587
seq001178
seq000545
seq000570
seq001054
seq000037
seq000081
seq001352
seq001385
seq000728
seq000103
seq000020
seq000039
seq001258
seq001925
seq000629
seq000671
seq000022
seq002765
seq000892
seq000354
seq003113
seq001029
seq000861
seq000503
seq000048
seq003023
seq000096
seq000754
seq003066
seq002840
seq001439
seq000649
seq001672
seq000443
seq000132
seq000138
seq001248
seq002264
seq001377
seq000600
seq000517
seq000854
seq001370
seq000722
seq002376
seq000206
seq003596
seq002210
seq003299
seq000104
seq000066
seq000617
seq000191
seq000767

seq000863
seq000504
seq000976
seq001017
seq001011
seq001066
seq000188
seq001042
seq000988
seq000833
seq000411
seq001461
seq000055
seq004387
seq000030
seq004166
seq001626
seq000834
seq003500
seq002743
seq000088
seq000297
seq001103
seq000055
seq002544
seq003066
seq000967
seq000034
seq000394
seq004799
seq000799
seq000378
seq001162
seq000378
seq001251
seq000452
seq001130
seq000345
seq002800
seq000603
seq000726
seq003510
seq000351
seq004799
seq000247
seq000619
seq000030
seq002425
seq001162
seq002080
seq001435
seq000582
seq001199
seq001521
seq000117
seq000519
seq003184
seq000405
seq000162
seq000035
seq000221
seq000174

seq000594
seq001677
seq000958
seq000070
seq001391
seq000313
seq000319
seq001532
seq001017
seq000054
seq000586
seq000633
seq004599
seq000930
seq001132
seq000184
seq000573
seq000434
seq005446
seq000087
seq001688
seq001698
seq003021
seq000877
seq000023
seq001069
seq000977
seq000291
seq000324
seq000649
seq008853
seq003113
seq000041
seq003681
seq004470
seq000909
seq001384
seq002259
seq001742
seq002507
seq003184
seq000026
seq002650
seq002530
seq000656
seq001190
seq008267
seq000027
seq000597
seq002343
seq000107
seq001660
seq002122
seq002430
seq004657
seq000137
seq001558
seq002840
seq000383
seq000506
seq000692
seq000307

seq000309
seq003023
seq000976
seq006079
seq000326
seq000805
seq000073
seq000400
seq001062
seq001519
seq002141
seq001102
seq000373
seq000111
seq000650
seq001487
seq003296
seq001149
seq000383
seq000607
seq000150
seq002079
seq004599
seq000442
seq000519
seq003352
seq002641
seq000470
seq000498
seq001208
seq000634
seq000653
seq004198
seq000299
seq002373
seq000298
seq001182
seq000042
seq003296
seq000603
seq000582
seq000159
seq005661
seq008180
seq000032
seq001352
seq000936
seq001237
seq000250
seq000537
seq003176
seq001636
seq000634
seq001093
seq000754
seq002617
seq000943
seq000816
seq000908
seq000087
seq001218
seq003093

seq001847
seq004392
seq001939
seq002507
seq001391
seq000716
seq002761
seq002091
seq003066
seq003520
seq000921
seq004944
seq000055
seq002111
seq002422
seq000344
seq001145
seq000874
seq000443
seq000792
seq001218
seq000581
seq001863
seq003500
seq002840
seq000554
seq001448
seq000966
seq003652
seq000247
seq001522
seq000406
seq001452
seq000555
seq000124
seq000208
seq000545
seq003901
seq002273
seq003184
seq000115
seq000790
seq013669
seq000875
seq002576
seq001178
seq000309
seq000570
seq003495
seq000887
seq004791
seq000664
seq005446
seq004420
seq000830
seq002554
seq003510
seq001062
seq001403
seq000014
seq002259
seq000978

seq000297
seq002162
seq004167
seq001435
seq000753
seq002415
seq001796
seq002045
seq000511
seq001591
seq001707
seq001847
seq001380
seq003067
seq005592
seq008537
seq000191
seq000615
seq000575
seq006304
seq000840
seq004392
seq000592
seq002882
seq000706
seq017366
seq003510
seq000344
seq000463
seq000884
seq000148
seq002784
seq001597
seq000022
seq000640
seq002201
seq001704
seq000976
seq004889
seq002249
seq000221
seq001108
seq000196
seq000231
seq002395
seq000615
seq000573
seq003611
seq001323
seq005557
seq001192
seq003431
seq001456
seq003169
seq001132
seq012358
seq004889
seq000176
seq000593
seq001617
seq002818
seq003196

seq002376
seq000172
seq003176
seq001674
seq004818
seq001276
seq008853
seq002821
seq000077
seq002201
seq001901
seq008853
seq009561
seq000736
seq002715
seq002288
seq000422
seq002149
seq001171
seq000738
seq000792
seq003021
seq001660
seq004898
seq003596
seq001452
seq001248
seq000592
seq001720
seq001412
seq000882
seq003196
seq004888
seq003520
seq005133
seq002595
seq000243
seq001464
seq001262
seq005592
seq000162
seq000089
seq001145
seq001224
seq001117
seq002576
seq001739
seq001059
seq000326
seq002363
seq002911
seq005131
seq000696
seq002881
seq002313
seq000422
seq001130
seq002091
seq000307
seq000661
seq000957
seq000146

seq000409
seq004387
seq000311
seq003979
seq000594
seq000470
seq000293
seq012487
seq000934
seq004392
seq001950
seq002280
seq003342
seq001446
seq002308
seq000592
seq001731
seq001856
seq000143
seq004541
seq004166
seq000302
seq002483
seq006930
seq004982
seq004599
seq002267
seq000325
seq002428
seq000690
seq000421
seq000364
seq000834
seq003162
seq000419
seq001066
seq000768
seq000108
seq001094
seq001322
seq001521
seq002764
seq002080
seq001038
seq000283
seq003848
seq001073
seq004641
seq019702
seq000876
seq005273
seq002816
seq002879
seq001689
seq000301
seq000463
seq001560
seq000190
seq000030
seq003258
seq001978
seq000889

seq005169
seq001147
seq003196
seq000921
seq000336
seq002390
seq002260
seq003299
seq003296
seq001127
seq003322
seq000287
seq002577
seq003597
seq000545
seq000089
seq003911
seq010124
seq000403
seq002744
seq001561
seq000795
seq000600
seq000134
seq000373
seq000622
seq004296
seq001431
seq000729
seq000436
seq000707
seq000604
seq001405
seq002363
seq012338
seq000016
seq000541
seq004538
seq000051
seq001196
seq002363
seq001522
seq000008
seq000595
seq005375
seq002743
seq001182
seq000642
seq002879
seq000085
seq004328
seq000137
seq001352
seq000078
seq000794
seq000780
seq002313
seq000291
seq000458
seq003611
seq004799
seq006342

seq000603
seq000506
seq000108
seq003595
seq000625
seq005446
seq000912
seq000129
seq004770
seq000105
seq000205
seq005592
seq003187
seq000650
seq001905
seq000142
seq000302
seq001037
seq003651
seq003559
seq006320
seq000912
seq000163
seq006780
seq003503
seq001925
seq000339
seq000515
seq000131
seq001014
seq002361
seq000198
seq003690
seq001042
seq001722
seq001347
seq012358
seq001219
seq002153
seq002507
seq000755
seq003632
seq002090
seq000409
seq001249
seq012161
seq002749
seq001507
seq000509
seq002870
seq001349
seq002764
seq004489
seq007357
seq000677
seq001572
seq019823
seq000316
seq002139
seq004192
seq002114
seq000584

seq002764
seq005103
seq010357
seq001145
seq002153
seq003355
seq000453
seq001117
seq001120
seq001183
seq000097
seq000925
seq002689
seq003192
seq003550
seq001993
seq000909
seq010289
seq000282
seq001196
seq001081
seq004051
seq004051
seq000854
seq002060
seq003399
seq001446
seq000210
seq014631
seq001231
seq000002
seq000439
seq000293
seq001770
seq003979
seq000984
seq015259
seq002645
seq005993
seq000365
seq000876
seq000378
seq003051
seq000039
seq003444
seq007705
seq001849
seq000406
seq000772
seq002264
seq005698
seq001259
seq009558
seq003067
seq000386
seq005408
seq000214
seq005531
seq000710
seq000101
seq001380
seq002395

seq003632
seq001228
seq002899
seq001827
seq002903
seq011063
seq004844
seq000567
seq001150
seq002618
seq004245
seq000111
seq000950
seq000015
seq002122
seq000072
seq000107
seq000595
seq005561
seq005133
seq000565
seq003246
seq001510
seq000293
seq000132
seq001387
seq001073
seq002832
seq004119
seq000093
seq003310
seq002267
seq001937
seq000247
seq000553
seq000642
seq001847
seq000560
seq000568
seq000323
seq000099
seq004944
seq003520
seq003604
seq000877
seq000876
seq006628
seq002323
seq004387
seq001453
seq001693
seq001066
seq000077
seq000194
seq000933
seq002940
seq017095
seq005698
seq002290
seq017046
seq002362
seq012358

seq000710
seq002461
seq002617
seq001439
seq000555
seq002158
seq000160
seq003985
seq000220
seq002098
seq011000
seq003023
seq002403
seq005560
seq002617
seq002737
seq006199
seq002838
seq004818
seq012570
seq001730
seq000408
seq000078
seq003302
seq002390
seq000542
seq000938
seq002272
seq000476
seq001675
seq009812
seq002689
seq001510
seq014354
seq001823
seq000811
seq000191
seq003071
seq004892
seq002800
seq003616
seq001452
seq000319
seq004462
seq003044
seq002821
seq000612
seq000575
seq000721
seq001764
seq004250
seq002056
seq008014
seq000697
seq002829
seq000103
seq012161
seq006609
seq004328
seq006304
seq000059
seq000615

seq002352
seq000642
seq000485
seq001363
seq000737
seq003729
seq004462
seq009619
seq000811
seq005718
seq006320
seq003375
seq000485
seq027227
seq006285
seq006022
seq000629
seq004107
seq003322
seq001786
seq003862
seq000569
seq005201
seq001238
seq003556
seq001626
seq000696
seq000282
seq002232
seq002045
seq000924
seq002313
seq003218
seq005103
seq001449
seq001028
seq006285
seq007380
seq000619
seq018528
seq000098
seq001901
seq002621
seq000240
seq003820
seq001553
seq000520
seq000636
seq000814
seq002749
seq001842
seq000250
seq001346
seq001797
seq010129
seq002522
seq000779
seq007087
seq000484

**Table S8 B -Significant OTUs identified by MaAsLin2 and ANCOM in NY**

| Ancom NY | MaAsLin2 NY | Common elements NY |
|---|---|---|
| seq000071 | seq000071 | seq000071 |
| seq000313 | seq000053 | seq000053 |
| seq000158 | seq000071 | seq000295 |
| seq000231 | seq000053 | seq000231 |
| seq000336 | seq000295 | seq000158 |
| seq000295 | seq000231 | seq000409 |
| seq000409 | seq000158 | seq000318 |
| seq000760 | seq000409 | seq000188 |
| seq000236 | seq000318 | seq000504 |
| seq000076 | seq000071 | seq000017 |
| seq000345 | seq000188 | seq000236 |
| seq001441 | seq000158 | seq000419 |
| seq000452 | seq000504 | seq000122 |
| seq000504 | seq000231 | seq000076 |
| seq000188 | seq000017 | seq000881 |
| seq000881 | seq000236 | seq000345 |
| seq000238 | seq000493 | seq000238 |
| seq000053 | seq000419 | seq000313 |
| seq000861 | seq000055 | seq001021 |
| seq000318 | seq001342 | seq000581 |
| seq000768 | seq000409 | seq000737 |
| seq000737 | seq000122 | seq000768 |
| seq000587 | seq000087 | seq000760 |
| seq001386 | seq000076 | seq000336 |
| seq001021 | seq000055 | seq000782 |
| seq000782 | seq000188 | seq000587 |
| seq000994 | seq000659 | |
| seq000017 | seq000881 | |
| seq000789 | seq000081 | |
| seq000122 | seq000017 | |
| seq001011 | seq000230 | |
| seq000581 | seq001423 | |
| seq001576 | seq000295 | |
| seq000706 | seq004744 | |
| seq001062 | seq000574 | |
| seq000419 | seq000318 | |
| | seq000019 | |
| | seq000345 | |
| | seq000236 | |
| | seq000770 | |
| | seq000238 | |
| | seq000574 | |
| | seq000138 | |
| | seq000041 | |
| | seq000146 | |
| | seq000320 | |
| | seq000189 | |
| | seq000345 | |
| | seq000645 | |
| | seq000081 | |
| | seq000087 | |
| | seq000055 | |
| | seq000409 | |
| | seq000726 | |
| | seq000462 | |
| | seq000389 | |
| | seq000001 | |
| | seq001766 | |
| | seq000477 | |
| | seq000813 | |
| | seq000813 | |
| | seq000750 | |
| | seq000003 | |
| | seq000122 | |
| | seq000419 | |
| | seq000711 | |
| | seq000023 | |
| | seq000053 | |

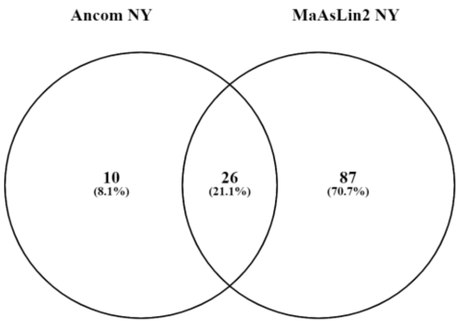

seq000313
seq000081
seq000077
seq001021
seq000019
seq000581
seq001342
seq000750
seq000504
seq001224
seq000737
seq000647
seq001342
seq000768
seq004744
seq000666
seq000760
seq000760
seq000016
seq000299
seq004847
seq000105
seq000073
seq000297
seq000297
seq001766
seq000029
seq000295
seq000336
seq000099
seq000782
seq000666
seq000702
seq004744
seq004847
seq000043
seq000732
seq003902
seq003881
seq003151
seq001639
seq007678
seq000228
seq000230
seq000384
seq001099
seq005959
seq000541
seq000428
seq000228
seq000402
seq000032
seq006932
seq000505
seq000058
seq001293
seq000310
seq001974
seq002181
seq001190
seq000395
seq000072
seq000152
seq000212
seq005926
seq000107
seq000311
seq000224
seq002005
seq009683

seq005418
seq000019
seq000740
seq000711
seq000693
seq008932
seq000772
seq000244
seq000853
seq003625
seq001353
seq000587
seq000085
seq000516
seq000132
